# The SARS-CoV-2 RNA interactome

**DOI:** 10.1101/2020.11.02.364497

**Authors:** Sungyul Lee, Young-suk Lee, Yeon Choi, Ahyeon Son, Youngran Park, Kyung-Min Lee, Jeesoo Kim, Jong-Seo Kim, V. Narry Kim

## Abstract

SARS-CoV-2 is an RNA virus whose success as a pathogen relies on its ability to repurpose host RNA-binding proteins (RBPs) to form its own RNA interactome. Here, we developed and applied a robust ribonucleoprotein capture protocol to uncover the SARS-CoV-2 RNA interactome. We report 109 host factors that directly bind to SARS-CoV-2 RNAs including general antiviral factors such as ZC3HAV1, TRIM25, and PARP12. Applying RNP capture on another coronavirus HCoV-OC43 revealed evolutionarily conserved interactions between viral RNAs and host proteins. Network and transcriptome analyses delineated antiviral RBPs stimulated by JAK-STAT signaling and proviral RBPs responsible for hijacking multiple steps of the mRNA life cycle. By knockdown experiments, we further found that these viral-RNA-interacting RBPs act against or in favor of SARS-CoV-2. Overall, this study provides a comprehensive list of RBPs regulating coronaviral replication and opens new avenues for therapeutic interventions.

## Introduction

Coronaviruses (CoVs) are a group of enveloped viruses with nonsegmented, single-stranded, positive-sense (+) RNA genomes, which belong to order *Nidovirales*, family *Coronaviridae*, and subfamily *Coronavirinae* (Lai and Cavanagh, 1997). They are classified into four genera: *Alphacoronavirus* and *Betacoronavirus* which exclusively infect mammals and *Gammacoronavirus* and *Deltacoronavirus* which primarily infect birds (Woo et al., 2012). Human CoVs such as *Alphacoronavirus* HCoV-229E and *Betacoronavirus* HCoV-OC43 have been known since the 1960s (Hamre and Procknow, 1966) as etiologic agents of the common cold. However, with the beginning of the 21st century, the world experienced the emergence of five novel human coronavirus species including highly pathogenic Severe acute respiratory syndrome coronavirus (SARS-CoV) in 2002 (Peiris et al., 2003), Middle East respiratory syndrome coronavirus (MERS-CoV) in 2012 (de Groot et al., 2013), and SARS-CoV-2 in 2019 (Zhou et al., 2020).

At the core of the coronavirus particle, the RNA genome is encapsulated in nucleocapsid (N) protein and surrounded by the viral membrane that contains spike (S) protein, membrane (M) protein, and envelope (E) protein (Lai and Cavanagh, 1997). The coronaviral RNA genome is ~30 kb which is the longest among RNA viruses and contains a 5’-cap structure and a 3’ poly(A) tail (Bouvet et al., 2010; Lai and Stohlman, 1981). Upon cell entry, the genomic RNA (gRNA) acts as an mRNA to produce nonstructural proteins (nsps) that are required for viral RNA production (Perlman and Netland, 2009). The ORF1a encodes polypeptide 1a (pp1a, 440-500 kDa) that is cleaved into 11 nsps. The −1 ribosomal frameshift occurs immediately upstream of the ORF1a stop codon, allowing translation of downstream ORF1b, yielding a large polypeptide (pp1ab, 740-810 kDa) that is cleaved into 15 nsps. Together, 16 different nsp fragments are generated to construct double-membrane vesicles (DMV) and mediate subsequent steps of viral RNA synthesis.

After this initial stage of viral translation, the gRNA is used as the template for the synthesis of negative-strand (-) RNA intermediates which in turn serve as the templates for positive-sense (+) RNA synthesis (Snijder et al., 2016; Sola et al., 2015). Ten different canonical (+) RNA species are produced from the SARS-CoV-2 genome, which include one full-length gRNA and nine subgenomic RNAs (sgRNAs) (Kim et al., 2020a). All canonical viral (+) RNAs share the common 5’ end sequence called the leader sequence and the 3’ end sequences. The sgRNAs are generated via discontinuous transcription which leads to the fusion between the 5’ leader sequence and the “body” parts containing the downstream open reading frames (Sola et al., 2015) that encode structural proteins (S, E, M, and N) and accessory proteins (3a, 3c, 6, 7a, 7b, 8, and 9b) (Kim et al., 2020a).

To accomplish this, coronaviruses employ unique strategies to evade, modulate, and utilize the host machinery (Fung and Liu, 2019). For example, the gRNA molecules must be kept in an intricate balance between translation, transcription, and encapsulation by recruiting the right host RNA-binding proteins (RBPs) and forming specific ribonucleoprotein (RNP) complexes. As host cells counteract by launching RBPs such as RIG-I, MDA5, and Toll-like receptors (TLRs) to recognize and eliminate viral RNAs, the virus needs to evade the immune system using its components to win the arms race between virus and host. How such stealthy devices are genetically coded in this compact RNA genome is yet to be explored (Snijder et al., 2016). Thus, the identification of the RBPs that bind to viral transcripts (or the SARS-CoV-2 RNA interactome) is key to uncovering the molecular rewiring of viral gene regulation and the activation of antiviral defense systems.

Biochemical techniques for studying RNA-protein interactions have been developed (Ramanathan et al., 2019) with the advancement in protein-centric methods such as CLIP-seq (crosslinking immunoprecipitation followed by sequencing) (Ule et al., 2018). In CLIP-seq experiments, RNP complexes are crosslinked by UV irradiation within cells to identify direct RNA-protein interactions. The protein of interest is immunoprecipitated to identify the associated RNAs (Lee and Ule, 2018; Van Nostrand et al., 2020). More recently, RNA-centric methods have also been developed to profile the mRNA interactome and RNP complexes (Roth and Diederichs, 2015). After UV irradiation, the RNA of interest is purified with oligonucleotide probes and the crosslinked proteins are identified by mass spectrometry. For example, RAP-MS exhibits compelling evidence of highly confident profiling of proteins that bind to a specific RNA owing to a combination of long hybridization probes and harsh denaturing condition (Engreitz et al., 2013; McHugh et al., 2015).

In this study, we developed a robust RNP capture protocol to define the repertoire of viral and host proteins that associate with the transcripts of coronaviruses, namely SARS-CoV-2 and HCoV-OC43. Network and transcriptome analyses combined with knockdown experiments revealed host factors that link the viral RNAs to mRNA regulators and putative antiviral factors.

## Results and Discussion

### SARS-CoV-2 RNP purification

To identify the viral and host proteins that directly interact with the genomic and subgenomic RNAs of SARS-CoV-2, we modified the RNA antisense purification coupled with mass spectrometry (RAP-MS) (McHugh and Guttman, 2018) protocol which was developed to profile the interacting proteins of a particular RNA species (Figure 1A). Briefly, cells were first detached from culture vessels and then irradiated with 254 nm UV to induce RNA-protein crosslink while preserving RNA integrity. Crosslinked cells were treated with DNase and lysed with an optimized buffer condition to homogenize and denature the proteins in high concentration. Massive pools of biotinylated antisense 90-nt probes were used to capture the denatured RNP complexes in a sequence-specific manner. After stringent washing and detergent removal, the RNP complexes were released and digested by serial benzonase and on-bead trypsin treatment. These modifications to the RAP-MS protocol enabled robust and sensitive identification of proteins directly bound to the RNA target of interest (see Methods for detailed explanation).

**Figure 1.**
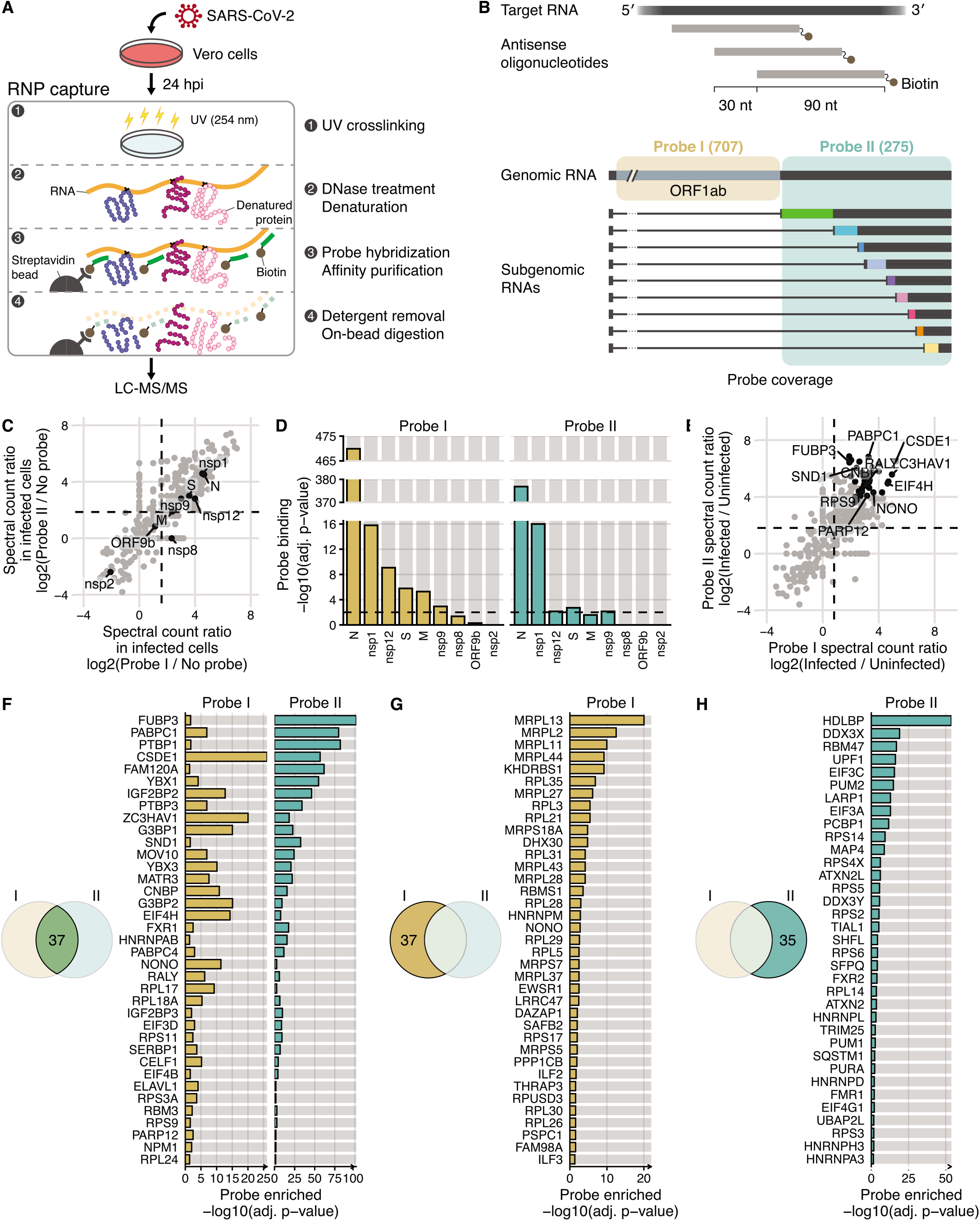
Comprehensive identification host and viral proteins that directly interact with the SARS-CoV-2 RNAs. (A) Schematic of the modified RAP-MS protocol in SARS-CoV-2-infected Vero cells. (B) Schematic of two separate pools of 90-nt antisense oligonucleotides and their SARS-CoV-2 RNA coverage. The first probe set “Probe I” consists of 707 oligonucleotides that cover the unique region of gRNA, and the second probe set “Probe II” consists of 275 oligonucleotides that cover the common region of gRNA and sgRNAs. (C) Spectral count ratio of Probe I (x-axis) and Probe II (y-axis) experiments over no-probe control in SARS-CoV-2-infected Vero cells (n = 3 technical replicates). Host proteins are marked by grey circles, and viral proteins (n = 9) are marked and labelled in black. The mean spectral count ratio of Probe I and of Probe II experiments are marked by vertical and horizontal dashed lines, respectively. (D) Statistical analysis of the quantity of viral proteins over no-probe control (i.e. probe binding). Adjusted p-values (adj. P-value) of Probe I experiments and of Probe II experiments are shown in yellow and green, respectively. Threshold for statistical significance (adj. p-value < 0.01) is indicated by horizontal dashed lines. (E) Spectral count ratio of Probe I (x-axis) and Probe II (y-axis) experiments in SARS-CoV-2-infected Vero cells compared to RNP captureexperiments in uninfected cells (n = 3 technical replicates). Statistically significant host proteins (n = 37, adj. P-value < 0.05) in both Probe I and Probe II experiments are marked by black circles. Of those, representative host proteins are labelled. The mean spectral count ratio of Probe I and of Probe II experiments are marked by vertical and horizontal dashed lines, respectively. (F) Statistical analysis of host proteins enriched in both Probe I and Probe II experiment (i.e. probe enriched). Adjusted p-values (adj. P-value) of Probe I experiments and of Probe II experiments are shown in yellow and green, respectively. (G) Statistical analysis of host proteins enriched in only Probe I experiment and (H) in only Probe II experiment.

We designed two separate pools of densely overlapping 90-nt antisense probes to achieve an unbiased perspective of the SARS-CoV-2 RNA interactome (Figure 1B and Table S1). The SARS-CoV-2 transcriptome consists of (1) a genomic RNA (gRNA) encoding 16 nonstructural proteins (nsps) and (2) multiple subgenomic RNAs (sgRNAs) that encode structural and accessory proteins (Sola et al., 2015). The sgRNAs are more abundant than the gRNA (Kim et al., 2020a). The first pool (“Probe I”) consists of 707 oligos tiles every 30 nucleotides across the ORF1ab region (266:21553, NC_045512.2) and thus hybridizes specifically with the gRNA molecules (Figure 1B). The second pool of 275 oligos (“Probe II”) covers the remaining region (21563:29872, NC_045512.2) which is shared by both the gRNA and sgRNAs.

To first check whether our method specifically captures the viral RNP complexes, we compared the resulting purification from Vero cells infected with SARS-CoV-2 (BetaCoV/Korea/KCDC03/2020) at MOI 0.1 for 24 hours (Kim et al., 2020b) by either Probe I or Probe II. As negative controls, we pulled-down without probes (“no probe” control) or with the control probes (for either 18S or 28S rRNA). Protein composition of each RNP sample was distinct as shown by silver staining and western blotting (Figure S1A) with prominent SARS-CoV-2 N protein associated with Probes I and II, as expected. Enrichment of SARS-CoV-2 RNAs were confirmed by RT-qPCR (Figure S1B), suggesting that our protocol purfies specific RNP complexes. Note that SARS-CoV-2 gRNA was not enriched in the Probe II experiment, hinting at the excess amount of sgRNAs over gRNA in our culture condition.

### Comprehensive identification of proteins binding to the SARS-CoV-2 transcripts

We conducted Label-free quantification (LFQ) by liquid chromatography with tandem mass spectrometry (LC-MS/MS) and identified 429 host proteins and 9 viral proteins in total (Figure 1C). As highly abundant proteins may nonspecifically co-precipitate during the RNP capture experiment, we statistically modelled this protein background as a multinomial distribution and assessed the probability (i.e. p-value) of the quantity of the identified protein in the RNP capture (e.g. Probe I) experiment over the protein background of the control (e.g. no-probe) experiment (see Methods for details). This unweighted spectral count analysis resulted in 199 and 220 proteins that are overrepresented in the Probe I and Probe II sample, respectively (FDR < 10%, Table S2). Protein domain enrichment analysis revealed that these proteins indeed harbor RNA-binding domains such as RNA recognition motif (RRM) domain and K Homology (KH) domain (Figure S1C). Of note, unlike the cellular mRNA interactome (Castello et al., 2012; Gerstberger et al., 2014), the RNA-binding repertoire of SARS-CoV-2 RNAs showed a depletion of DEAD/DEAH box helicase domains and an enrichment of KH domain.

As for viral proteins, the N protein was the most strongly enriched one, as expected (Figure 1D). The nsp1 protein was also statistically enriched in both Probe I and Probe II experiments. Nsp12, S, M, and nsp9 were detected more with Probe I than with Probe II. Coronavirus nsp9 is a single-strand RNA binding protein (Egloff et al., 2004; Sutton et al., 2004) essential for viral replication (Miknis et al., 2009). Nsp1 is one of the major virulence factors that suppresses host translation by binding to the 40S ribosomal subunit (Thoms et al., 2020). While nsp1 is mostly studied in the context of host gene expression (Narayanan et al., 2008), our result hints at the direct role of nsp1 on the transcripts of SARS-CoV-2.

To delineate the host proteins that are enriched in the SARS-CoV-2 RNP complex, we employed an additional negative control experiment with uninfected cells (Figure 1E). In effect, this control provides a conservative background of host proteins as shown by silver staining (Figure S1D). Distributions of peptide length were consistent across technical replicates (Figure S1E), demonstrating the robustness of the “on-bead” digestion step. Spectral count analysis against the “uninfected probe control” resulted in 74 and 72 proteins that are enriched in the infected samples with Probe I and Probe II, respectively (FDR < 5%, Figure 1F-H). In combination, we define these 109 proteins as the “SARS-CoV-2 RNA interactome.” 37 host proteins such as CSDE1 (Unr), EIF4H, FUBP3, G3BP2, PABPC1, ZC3HAV1 were enriched in both the Probe I and Probe II RNP capture experiments on infected cells (Figure 1F), thus identifying a robust set of the “core SARS-CoV-2 RNA interactome.” Gene ontology (GO) term enrichment analysis revealed that these host factors are involved in RNA stability control, mRNA function, and viral process (Figure S1F).

### Conservation of betacoronavirus RNP complexes

To investigate the evolutionary conservation of the RNA-protein interactions in coronaviruses, we conducted RNP capture on HCoV-OC43 that belongs to the lineage A of genus *betacoronavirus.* HCoV-OC43 shows 54.2% nucleotide homology to SARS-CoV-2 which belongs to lineage B. We profiled the HCoV-OC43 RNP complexes at multiple time points: 12, 24, 36, and 48 hours post-infection (Figure 2A, S2A and S2B). Two antisense probe sets (i.e. OC43 Probe I and OC43 Probe II) were designed in a similar manner as in SARS-CoV-2 experiments. OC43 Probe I hybridizes only with the gRNA of HCoV-OC43, while Probe II captures both gRNA and sgRNA molecules. As negative controls, the no-probe control on infected cells and the probe control on uninfected cells were used and confirmed by silver staining (Figure S2C).

**Figure 2.**
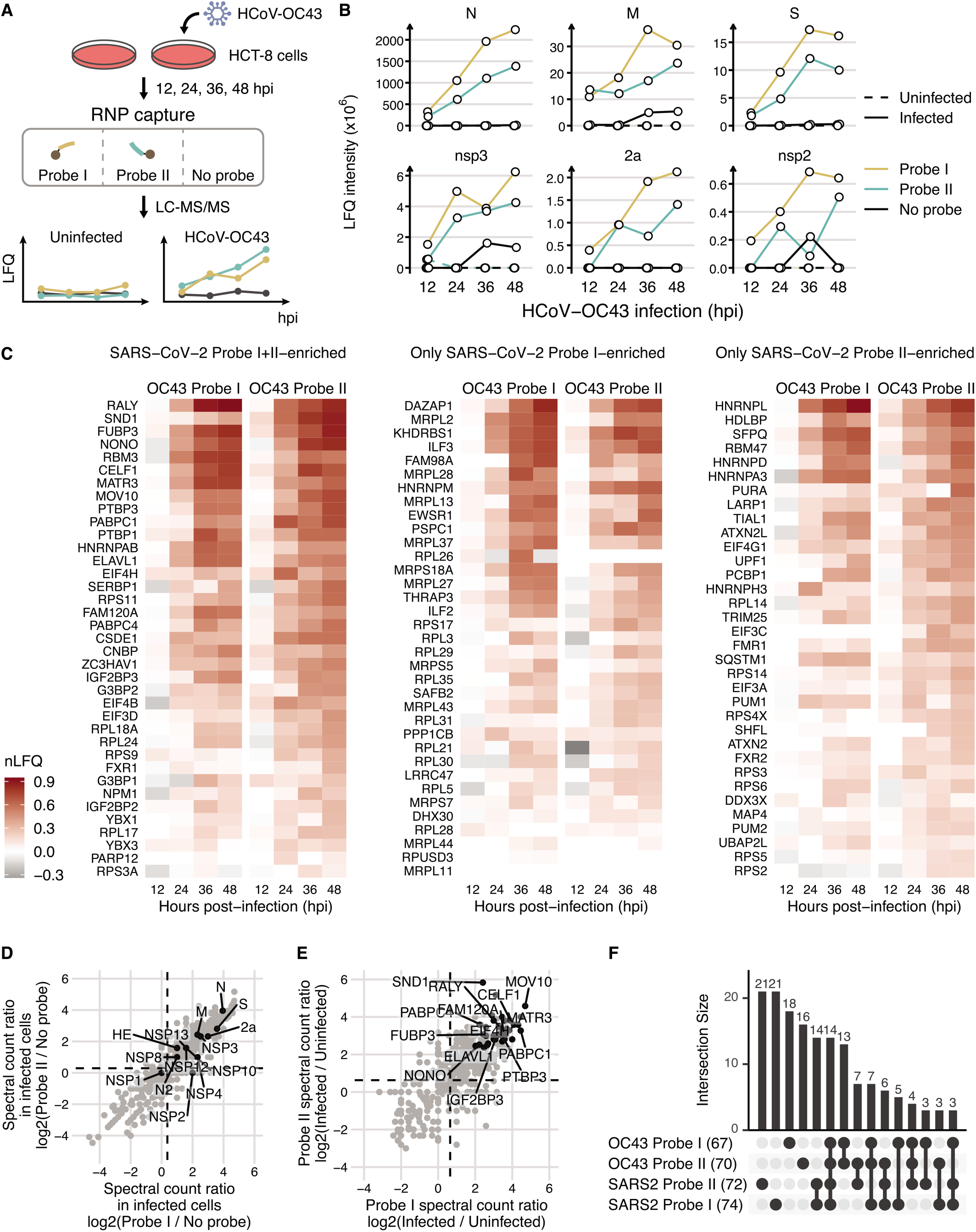
Comparison of SARS-CoV-2 and HCoV-OC43 RNA interactome. (A) Schematic of time-course RNP capture experiment in HCoV-OC43-infected HCT-8 cells. (B) LFQ intensity of abundant viral proteins identified in HCoV-OC43 RNP capture experiment at 12, 24, 36, and 48 hours post-infection (hpi). (C) Heatmap of normalized LFQ intensity (nLFQ) of HCoV-OC43 experiment of host proteins enriched in (left) both SARS-CoV-2 Probe I and Probe II experiments, (middle) only SARS-CoV-2 Probe I experiment, and (right) only SARS-CoV-2 Probe II experiment. nLFQ is the log10 fold change over the median LFQ intensity across each probe set (i.e. Probe I or Probe II). A pseudo-value of 1 ⍰ 10 was added to handle missing values. (D) Spectral count ratio of Probe I (x-axis) and Probe II (y-axis) experiments over no-probe control in HCoV-OC43-infected HCT-8 cells of 36 hpi. Host proteins are marked by grey circles, and viral proteins (n = 14) are marked and labelled in black. The mean spectral count ratio of Probe I and of Probe II experiments are marked by vertical and horizontal dashed lines, respectively. (E) Spectral count ratio of Probe I (x-axis) and Probe II (y-axis) experiments in HCoV-OC43-infected HCT-8 cells compared to uninfected cells of 36 hpi. Statistically significant host proteins (n = 38, adj. P-value < 0.05) in both Probe I and Probe II experiments are marked by black circles. Of those, representative host proteins are labelled. The mean spectral count ratio of Probe I and of Probe II experiments are marked by vertical and horizontal dashed lines, respectively. (F) Upset plot of HCoV-OC43 Probe I-enriched (n = 67), HCoV-OC43 Probe II-enriched (n = 70), SARS-CoV-2 Probe I-enriched (n = 73), and SARS-CoV-2 Probe II-enriched (n = 72) proteins. 14 host proteins were enriched in all four experiments.

Unweighted spectral count analysis against the no-probe control revealed 133, 167, 192, and 160 proteins that are overrepresented in the OC43 Probe I experiment at 12, 24, 36, and 48 hours post-infection (hpi), respectively (FDR < 10%, Table S3). For the OC43 Probe II experiment, 119, 189, 194, and 185 proteins were overrepresented at each respective infection time point of 12, 24, 36, and 48 hpi (FDR < 10%, Table S4). The analysis of all eight RNP capture experiments resulted in the enrichment of proteins containing canonical RNA binding domains such as the RRM domain and the KH domain (Figure S2D), indicating the specificity of the coprecipitated RBPs.

Fourteen viral proteins were detected within the HCoV-OC43 RNP complexes (Figure 2B, Figure S2E). Specifically, LFQ intensities for viral structural proteins N, M, and S increased over time more evidently in the gRNA-hybridizing Probe I experiment (Figure 2B). HCoV-OC43 2a, an accessory protein unique to *betacoronavirus* lineage A, was also detected and increased over time, indicating that this protein of unknown function may act as an RBP. The RdRP nsp12 and the papain-like protease nsp3 also increased along with the other nsps identified in this experiment. Only a marginal amount of the HCoV-OC43 nsp1 was detected (Figure S2E), implying the functional divergence of nsp1 in *betacoronavirus* lineages A and B.

Next, we compared the host factors that form the viral RNA interactome of HCoV-OC43 and SARS-CoV-2. All 107 proteins from the SARS-CoV-2 interactome were also detected in the HCoV-OC43 interactome throughout multiple infection timepoints, except for RBMS1 and DDX3Y (Figure 2C), indicating that many of the same host proteins interact with RNAs of both HCoV-OC43 and SARS-CoV-2. To determine the core host factors that are conserved in the coronavirus RNA interactomes, we applied our spectral count analysis on the HCoV-OC43 experiment of 36 hpi (Figure S2A) and conducted statistical analysis in comparison to the no-probe control (Figure 2D, FDR < 10%) and the uninfected-probe control (Figure 2E, FDR < 5%). We identified 67 and 70 host proteins for the HCoV-OC43 Probe I and Probe II experiments, respectively (Table S5). 38 proteins were statistically enriched in both probe sets. GO term enrichment analysis revealed that these 38 host proteins are involved in transcriptional regulation, RNA processing, and RNA stability control (Figure S2F). Of the 38 common proteins, 14 host proteins (CELF1, EIF4H, ELAVL1, FAM120A, FUBP3, IGF2BP3, MATR3, MOV10, NONO, PABPC1, PABPC4, PTBP3, RALY, SND1 and ZC3HAV1) were also found in the SARS-CoV-2 experiments (Figure 2F), and thus are conserved host components of the betacoronavirus RNA interactome.

### Regulatory landscape of the SARS-CoV-2 RNA interactome

To understand the regulatory significance of the SARS-CoV-2 RNA interactome, we compiled a list of “neighboring” proteins that are known to physically interact with the factors identified in our study (see Methods for details). In particular, we generated a physical interaction network centered (or seeded) by the core SARS-CoV-2 interactome (Figure S3A). Network analysis revealed several network hubs (e.g. NPM1 and PABPC1) and two highly connected network modules: the ribosomal subunits and the EIF3 complex. GO term enrichment analysis resulted in translation-related biological processes (Figure S3B), most likely due to the overrepresentation of ribosomal proteins and subunits of the EIF3 complex, which reflects the active translational status of viral mRNPs.

To achieve a more indepth functional perspective of the RNA interactome, we reconstructed the physical interaction network with the SARS-CoV-2 RNA interactome but excluding ribosomal proteins and EIF3 proteins (Figure 3A). This analysis identified additional hub proteins such as TRIM25, SQSTM1, and KHDRBS1. GO term enrichment analysis revealed multiple steps of the mRNA life cycle such as mRNA splicing, mRNA export, mRNA stability, and stress granule assembly (Figure 3B), suggesting these mRNA regulators are co-opted to assist the viral life cycle. Interestingly, we also found GO terms related to viral processes and innate immune response. In terms of intracellular localization, the SARS-CoV-2 RNA interactome is enriched by proteins localized in the paraspeckle and cytoplasmic RNP granule (e.g. stress granule) compared to the cellular mRNA interactome (Figure 3C) (Baltz et al., 2012; Castello et al., 2012). These observations suggest the regulatory mechanisms of viral RNAs distinct from that of host mRNAs, which involve activation of host antiviral machinery and sequestration of viral RNAs.

**Figure 3.**
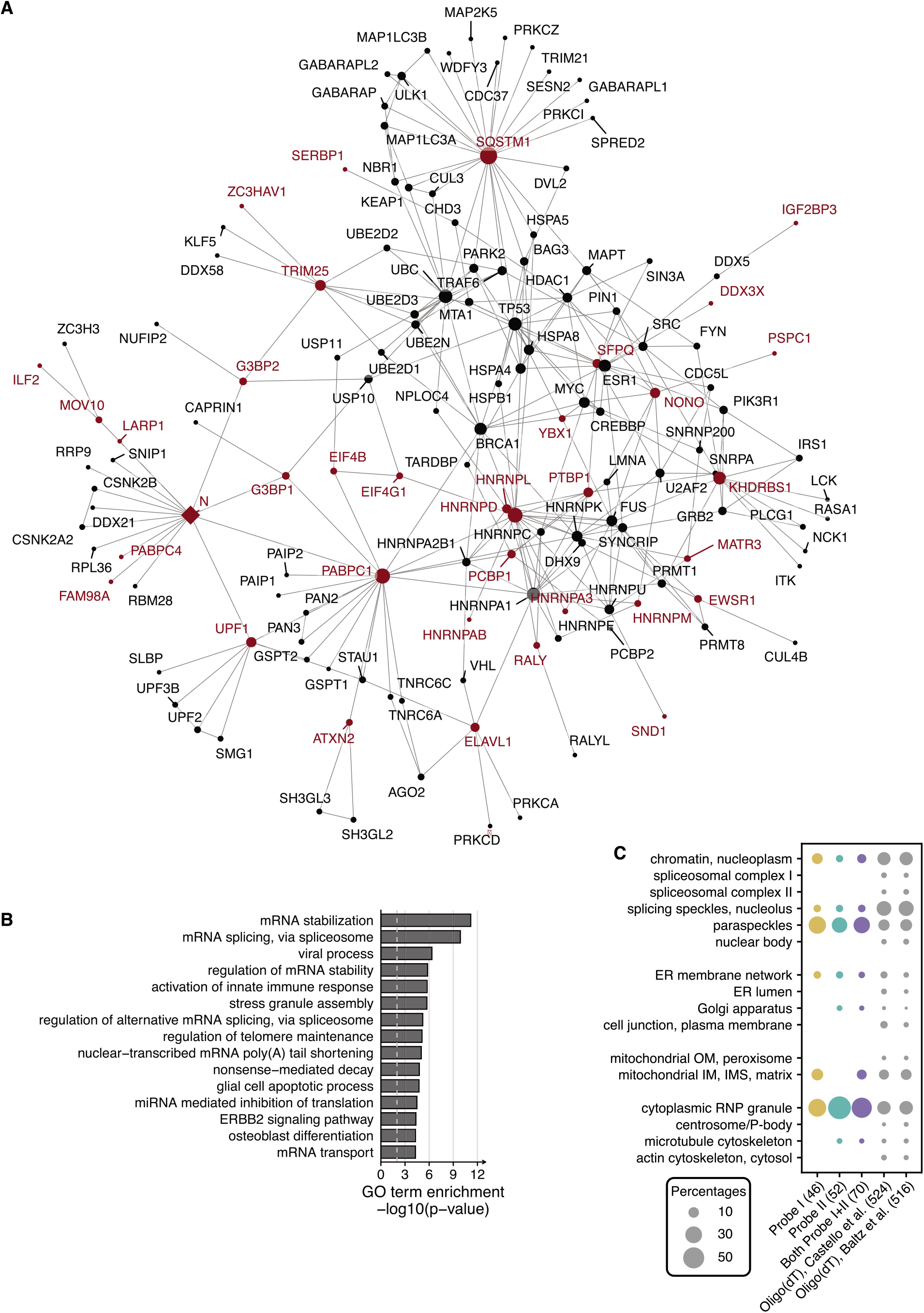
Molecular network of the SARS-CoV-2 RNA interactome. (A) Physical-interaction network of SARS-CoV-2 RNA interactome but excluding ribosomal proteins and EIF3 proteins. Host factors are marked in red. Host and viral proteins are marked by circles and diamonds, respectively. The size of the network node is proportional to the number of network edges (i.e. degree) of the node. Largest connected component of the network was chosen for visualization and interpretation. (B) Gene Ontology Biological Process term enrichment analysis of proteins retrieved in (A). (C) Subcellular localization of the SARS-CoV-2 RNA interactome and cellular mRNA interactomes. The number of proteins with localization information is shown in parentheses.

Another way to gauge the regulatory potential of the SARS-CoV-2 RNA interactome is to examine them in the context of the transcriptional response to viral infection. For example, infected cells recognize the non-cellular RNAs by a number of cytosolic sensors such as RIG-I (DDX58) and MDA5 (IFIH1) and ultimately induces interferons which in turn up-regulates interferon-stimulated genes (ISGs) through the JAK-STAT pathway (Sa Ribero et al., 2020). Multiple studies have reported the unusually low-level of type I/III interferon responses in cell and animal model systems of SARS-CoV-2 infection (Blanco-Melo et al., 2020) and blood biopsy from COVID-19 patients (Hadjadj et al., 2020), indicating active immune evasion by SARS-CoV-2 and supporting the therapeutic potential of timely interferon treatment (Sa Ribero et al., 2020).

To investigate the regulation of the SARS-CoV-2 RNA interactome, we utilized published transcriptome data of SARS-CoV-2-infected cells (Blanco-Melo et al., 2020). Transcriptome analysis of ACE2-expressing A549 cells revealed host factors of SARS-CoV-2 RNA interactome that are differentially expressed after infection (Figure 4A). Specifically, PARP12, SHFL, CELF1, and TRIM25 are up-regulated upon infection. Treatment of ruxolitinib, a JAK1 and 2 inhibitor, in infected cells suppressed the expression of 5 host factors (MOV10, PARP12, SHFL, TRIM25, and ZC3HAV1) (Figure 4B). TRIM25 and PARP12 are part of the 62 vertebrate core ISGs (Shaw et al., 2017). Interferon-beta (INFß) treatment on normal human bronchial epithelial (NHBE) cells induces PARP12, SHFL, and TRIM25 (Figure 4C). Consistently, proteome analysis of INFß-treated cells (Kerr et al., 2020) exhibited an upregulation of PARP12, TRIM25, and ZC3HAV1 (Figure S3C), altogether indicating that JAK-STAT signaling pathway regulates part of the SARS-CoV-2 RNA interactome. Moreover, reanalysis of mRNA-seq data of postmortem lung samples from COVID-19 patients (Blanco-Melo et al., 2020) showed up-regulation of SHFL and ZC3HAV1 (Figure 4D), highlighting the physiological relevance of SARS-CoV-2 RNP regulation in natural human infection of SARS-CoV-2.

**Figure 4.**
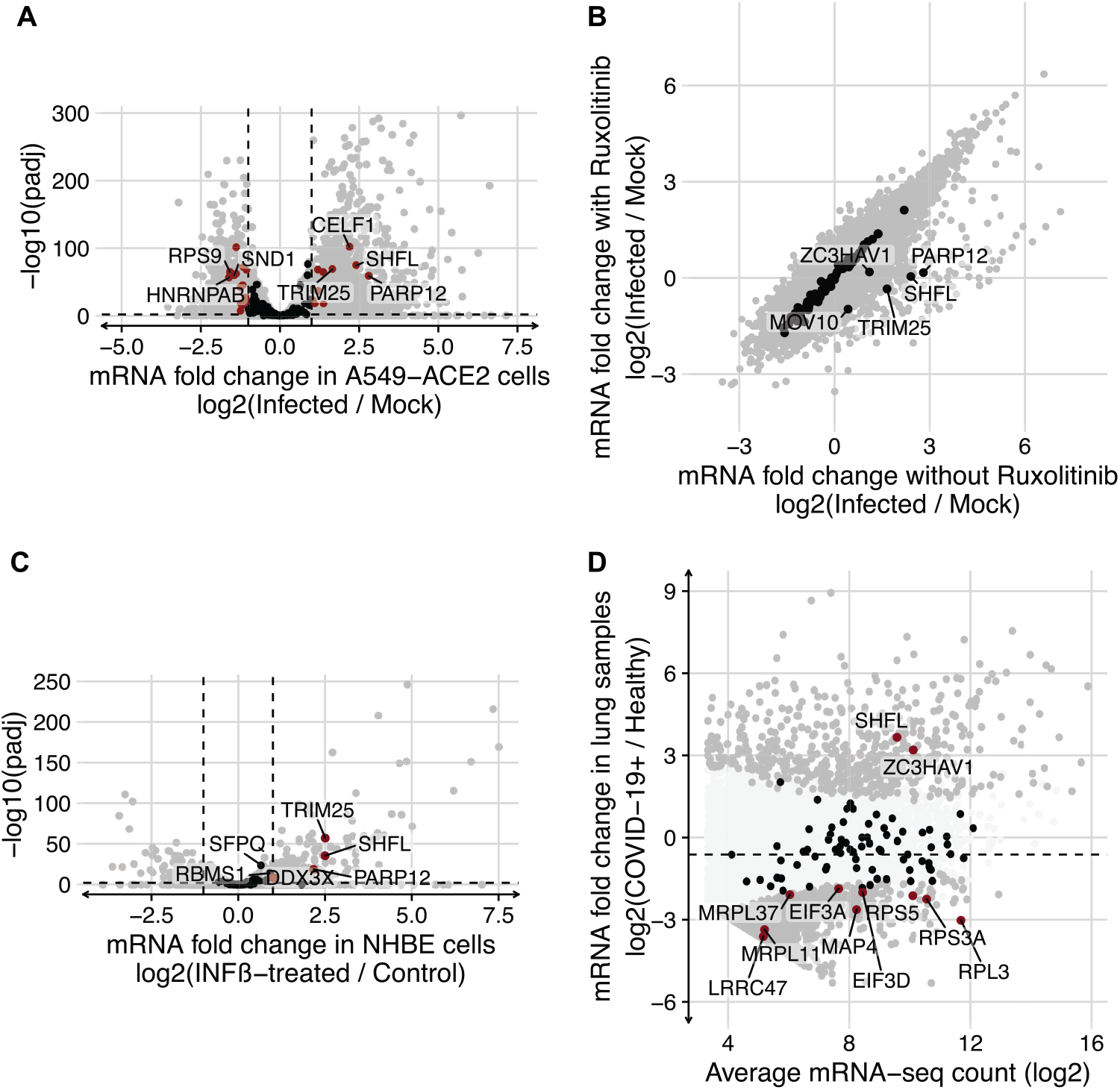
Regulation of the SARS-CoV-2 RNA interactome. (A) Differential expression analysis of ACE2-expressing A549 cells (A549-ACE2) after SARS-CoV-2 infection (MOI = 2.0). mRNA fold change (FC) of −1 and 1 are marked by vertical dashed lines. 1% FDR is marked by a horizontal dashed line. Host genes of the SARS-CoV-2 RNA interactome are marked in red if differentially expressed (FDR < 1% and |FC| > 1) or otherwise in black. (B) Comparison of mRNA fold change of SARS-CoV-2-infected (MOI = 2.0) A549-ACE2 cells after treatment of ruxolitinib. Host genes of the SARS-CoV-2 RNA interactome are marked in black. (C) Differential expression analysis of normal human bronchial epithelial (NHBE) cells after INFß treatment. Graph notations as in (D). (D) MA plot of published mRNA-seq data of post-mortem lung samples from COVID-19 patients and healthy lung tissue from uninfected individuals. Mean fold change is marked by a horizontal dashed line. Host genes of the SARS-CoV-2 RNA interactome are marked in red if differentially expressed (FDR < 5%) or otherwise in black. Other differentially expressed genes are in dark grey.

### Host factors and functional modules that regulate the SARS-CoV-2 RNAs

To measure the impact of these host proteins on coronavirus RNAs, we conducted knockdown experiments and infected Calu-3 cells with SARS-CoV-2 (Figure 5A and 5B). Calu-3 cells are human lung epithelial cells and often used as a model system for coronavirus infection (Sims et al., 2008). Strategically, we selected a subset of the SARS-CoV-2 RNA interactome that covers a broad range of functional modules that we identified above: JAK-STAT signaling, mRNA transport, mRNA stability, and translation. Knockdown of host factors that are stimulated by SARS-CoV-2 infection or INFß treatment, namely PARP12, TRIM25, ZC3HAV1, CELF1, and SHFL, led to increased viral RNAs (Figure 5C). The result suggests that these RBPs which directly recognize the viral RNAs are induced by the interferon and JAK-STAT signaling pathway to suppress coronaviruses.

**Figure 5.**
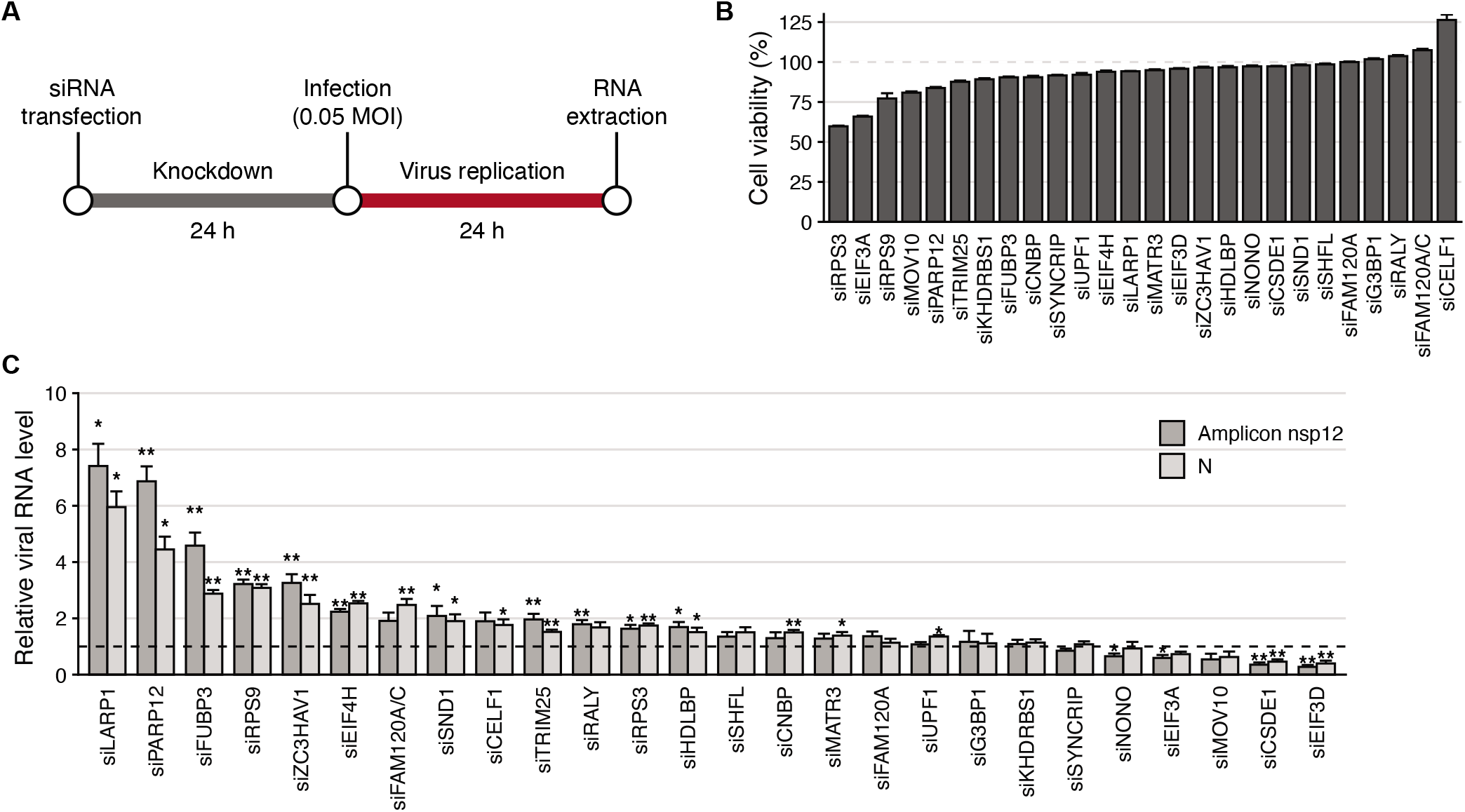
Effects of RBP depletion on viral RNA abundance. (A) Schematic of siRNA knockdown in SARS-CoV-2-infected Calu-3 cells. (B) Cell viability of siRNA-treated cells measured by MTT assay. Data are represented as mean ± s.e.m. (n□=□3 independent experiments). siNC was used as negative control. (C) Change in gRNA (amplicon nsp12, dark grey) and N sgRNA (amplicon N, light grey) levels after siRNA depletion were measured by RT-qPCR. Data are represented as mean ± s.e.m. (n□=□3 or 6 independent experiments). RNA levels in siNC-treated cells were used as negative control and marked by a horizontal dashed line. 18S rRNA was used for normalization. * P□<<#x25A1;0.05, ** P□<□Q.01; two-sided Student’s t-test.

ZC3HAV1 (ZAP/PARP13) is an ISG and known to restrict the replication of many RNA viruses such as HIV-1 (*Retroviridae*), sindbis virus (*Togaviridae*), and Ebola (*Filoviridae*) (Goodier et al., 2015). ZC3HAV1 was previously reported to recognize CpG and recruits decay factors to degrade HIV RNAs (Takata et al., 2017). ZC3HAV1 acts as a cofactor for TRIM25, an E3 ubiquitin ligase that promotes antiviral signaling mainly through RIG-I (Choudhury et al., 2020; Gack et al., 2007). Our knockdown results indicate that both ZC3HAV1 and TRIM25 may act as antiviral factors against SARS-CoV-2 (Figure 5C). SARS-CoV N protein was previously shown to interact with the SPRY domain of TRIM25, interfering with the activation of RIG-I (Hu et al., 2017). Further investigation is needed to understand how ZC3HAV1 and TRIM25 recognize and suppress SARS-CoV-2 transcripts and if SARS-COV-2 N protein counteracts TRIM25.

PARP12, a cytoplasmic mono-ADP-ribosylation (MARylation) enzyme, is also known to have broad antiviral activity against RNA viruses such as Venezuelan equine encephalitis virus (*Togaviridae*), vesicular stomatitis virus (*Rhabdoviridae*), Rift Valley fever virus (*Phenuiviridae*), encephalomyocarditis virus (*Picornaviridae*), and Zika virus (*Flaviviridae*) by multiple mechanisms including blocking cellular RNA translation (Atasheva et al., 2014; Welsby et al., 2014) or triggering proteasome-mediated destabilization of viral proteins (Li et al., 2018). ADP ribosyltransferases are evolutionarily ancient tools used for host-pathogen interactions (Fehr et al., 2020). Of note, coronavirus nsp3 carries a conserved macrodomain that can remove ADP-ribose to reverse the activity of PARP enzymes (Fehr et al., 2015). Knockdown of PARP12 and PARP14 was shown to increase the replication of the macrodomain-deficient mouse hepatitis virus (MHV) which belongs to the lineage A of genus *betacoronavirus* (Grunewald et al., 2019) which is consistent with our knockdown results (Figure 5C). Based on our RNA interactome data, we hypothesize that the RNA-binding activity of PARP12 and its role in viral RNA recognition may explain the underlying molecular mechanism of its antiviral activity against SARS-CoV-2 transcripts.

Other interferon-stimulated RBPs may also be involved in host defense against SARS-CoV-2. CELF1 (CUGBP1 Elav-like protein family 1) mediates alternative splicing and controls mRNA stability and translation (Konieczny et al., 2014). CELF1 is required for IFN-mediated suppression of simian immunodeficiency virus (*Retroviridae*) (Dudaronek et al., 2007), but its involvement in other viral infections is unknown. SHFL (Shiftless/RyDEN) was induced prominently upon viral infection and interferon treatment, and suppressed by JAK inhibitor (Figure 4). SHFL suppresses the translation of diverse RNA viruses, including dengue virus (*Flaviviridae*) and HIV (*Retroviridae*) (Balinsky et al., 2017; Suzuki et al., 2016; Wang et al., 2019). Under our experimental condition, upregulation of viral RNA was only modest in CELF1- and SHFL-depleted cells, but further examination is needed as the knockdown efficiency of ISGs were low in infected cells (Figure S4B), likely because virus-induced interferon response compromised gene silencing.

Apart from the above RBPs, we identified multiple host factors that have not been previously described in the context of viral infection. In particular, LARP1 depletion resulted in a substantial upregulation of viral RNAs (Figure 5C), suggesting that LARP1 may have an antiviral function. LARP1 stabilizes 5’ TOP mRNAs encoding ribosomal proteins and translation factors, which contain the 5’ terminal oligopyrimidine (5’ TOP) motif in the 5’ UTR (Philippe et al., 2018). LARP1 also represses the translation of 5’ TOP mRNAs in response to metabolic stress in an mTORC1-dependent manner (Hong et al., 2017; Lahr et al., 2017). While much is unknown regarding the role of LARP1 during viral infection, a recent proteomics study showed that SARS-CoV-2 N protein binds to LARP1 (Gordon et al., 2020). Based on our RNP capture and knockdown results, it is conceivable that LARP1 may interfere with viral translation directly via viral RNA interaction. The role of N protein in the context of LARP1-dependent viral suppression warrants further investigation. Of note, we cannot exclude the possibility that LARP1 indirectly regulates SARS-CoV-2 RNAs via the control of 5’ TOP mRNAs encoding translation machinery.

In fact, the SARS-CoV-2 RNA interactome includes specific components of the 40S and 60S ribosomal subunits and translational initiation factors (Figure 1F-H). Knockdown experiments indicated that ribosomal proteins (RPS9 and RPS3) and translation initiation factor EIF4H may have antiviral activities (Figure 5C). EIF4H along with EIF4B is a cofactor for RNA helicase EIF4A (Rogers et al., 2001) whose depletion results in RNA granule formation (Tauber et al., 2020). EIF4H and EIF4B were both identified as the core SARS-CoV-2 RNA interactome (Figure 1F). EIF4H also interacts with SARS-CoV-2 nsp9 (Gordon et al., 2020), so it will be interesting to investigate the functional consequence of the EIF4H-nsp9 interaction. Together, our observations implicate that SARS-CoV-2 infection may be closely intertwined with the regulation of ribosome biogenesis, metabolic rewiring, and global translational control.

Other translation factors EIF3A, EIF3D, and CSDE1 exhibited proviral effects (Figure 5C). EIF3A is the RNA-binding component of the mammalian EIF3 complex and evolutionarily conserved along with EIF3B and EIF3C (Masutani et al., 2007). EIF3D is known to interact with mRNA cap and is required for specialized translation initiation (Lee et al., 2016). CSDE1 (Unr) is required for IRES-dependent translation in human rhinovirus (*Picornaviridae*) and poliovirus (*Picornaviridae*) (Anderson et al., 2007; Boussadia et al., 2003). In all, our finding suggests that SARS-CoV-2 may recruit EIF3D and CSDE1 to respectively regulate cap-dependent and IRES-dependent translation initiation (Lee et al., 2017) of SARS-CoV-2 gRNA and sgRNAs.

Lastly, the coronaviral RNA interactomes are enriched with RBPs with KH domains (Figure S1D and Figure S2B) unlike the mRNA interactome. Depletion of FUBP3 (MARTA2) and HDLBP (Vigilin) increased the viral RNA levels (Figure 5C), hinting at a potential antiviral role of proteins containing KH domains. HDLBP is a conserved protein that contains 14 KH domains and has been implicated in viral translation of dengue virus (Ooi et al., 2019). FUBP3 was enriched in all four RNP capture experiments (i.e. SARS-CoV-2 Probe I/II and OC43 Probe I/II) (Figure 2D). FUBP3 is a nuclear protein with 4 KH domains and binds to the 3’ UTR of cellular mRNAs regulating mRNA localization (Blichenberg et al., 1999; Mukherjee et al., 2019). Its connection to the life cycle of coronavirus is unknown to our knowledge.

Our current study reveals a broad-spectrum of antiviral factors such as TRIM25, ZC3HAV1, PARP12, and SHFL and also many RBPs whose functions are unknown in the context of viral infection such as LARP1, FUBP3, FAM120A/C, EIF4H, RPS3, RPS9, SND1, CELF1, RALY, CNBP, EIF3A, EIF3D, and CSDE1. This list of proteins reflects constant host-pathogen interactions and opens new avenues to explore unknown mechanisms of viral life cycle and immune evasion.

Along with proteins regulating RNAs, it would also be interesting to consider the possibility of ‘riboregulation’ (Hentze et al., 2018) in which RNA controls its interacting proteins. Dengue virus, for example, uses its subgenomic RNA called sfRNA to sequester TRIM25 (Chapman et al., 2014). The sgRNA/gRNA ratio is a critical determinant of epidemic potential of dengue virus (Manokaran et al., 2015). Notably, coronaviruses including SARS-CoV-2 produces substantial amounts of noncanonical sgRNAs that may serve as noncoding decoys to interact with host RBPs to modulate host immune responses (Kim et al., 2020a).

There are over 3,200 human clinical studies listed for the treatment of COVID-19 as of September 2020 (ClinicalTrials.gov), yet there’s no effective antiviral drug or vaccine available to stop the ongoing pandemic. This unmet medical need highlights our substantial lack of knowledge of SARS-CoV-2. Thus, redefining antiviral strategies should be contemplated beyond expeditious drug repurposing efforts. So far, collective large and multidisciplinary datasets from viral transcriptome (Kim et al., 2020a), host transcriptional response (Blanco-Melo et al., 2020), ribosome profiling (Finkel et al., 2020), whole proteomics (Bojkova et al., 2020), protein-protein interactions by co-IP (Gordon et al., 2020) and proximity labeling (St-Germain et al., 2020), phosphoproteomics (Bouhaddou et al., 2020), RNA structure (Lan et al., 2020), genome-wide CRISPR screen (Wei et al., 2020), and off-label drug screening (Chen et al., 2020) have all provided invaluable insights of the underlying biology of this novel human coronavirus.

In line with these efforts, we here present the SARS-CoV-2 RNA interactome that offers insights into the host-viral interaction that regulate the life cycle of coronaviruses. Data interpretation in the context of publicly available orthogonal information has enabled the identification of proviral and antiviral protein candidates. We expect that further efforts to generate and integrate system-level data will elucidate the pathogenicity of SARS-CoV-2 and introduce new strategies to combat COVID-19.

## ACKNOWLEDGMENTS

Authors thank Sun-Je Woo for the support in BL3 laboratory, Kwangmi Moon for the advice on BL2 virus culture, and Soohyun Jang for technical support. We also thank Dr. Kwangseog Ahn for sharing reagents. This work would not have been possible without the invaluable discussions from members of Narry Kim’s lab, particularly Dongwan Kim, Haedong Kim, and Dr. Hyunjoon Kim. The pathogen resource for this study was provided by the National Culture Collection for Pathogens, Korea National Institute of Health for SARS-CoV-2 (NCCP43326) and Korea Bank for Pathogenic Viruses, Korea University Medical Center for HCoV-OC43 (KBPV-VR-8). This work was supported by the Institute for Basic Science from the Ministry of Science and ICT of Korea (IBS-R008-D1 to V.N.K.) and BK21 Research Fellowship from the Ministry of Education of Korea (A.S.).

## AUTHOR CONTRIBUTIONS

Conceptualization, S.L., Y.L., and V.N.K.; Methodology and experiments, S.L., A.S., and Y.P.; LC-MS/MS: J.K. and J-S.K.; Data analysis, Y.L. and Y.C; Manuscript writing, S.L., Y.L., and V.N.K.; Visualization, Y.L, Y.C., and A.S.; Supervision, V.N.K.

## Supplementary Figure Legends

**Figure S1.**
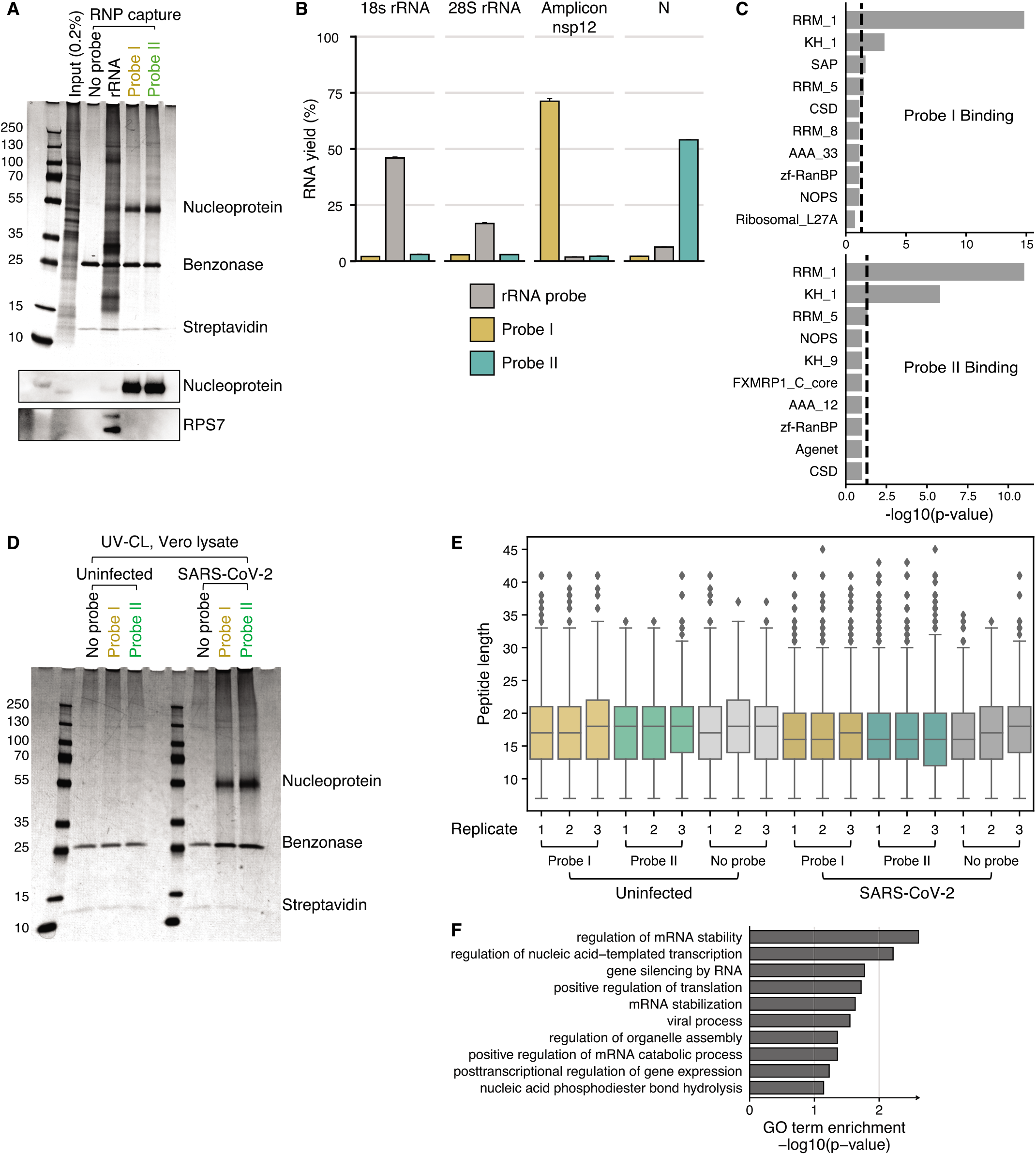
Experimental and statistical validation of modified RAP-MS experiments on SARS-CoV-2 RNAs. (A) Validation of antisense oligo-based RNA-protein (RNP) capture by silver staining and western blotting in SARS-CoV-2-infected Vero cells. SARS-CoV-2 nucleocapsid and RPS7 were used as control for rRNA and SARS-CoV-2 RNA (Probe I and Probe II) RNP capture experiments. CL: cross-linked. (B) RNA yield of antisense oligo-based RNA-protein (RNP) capture (rRNA, Probe I, and Probe II) measured by RT-qPCR (n = 2 technical replicates). Note that the SARS-CoV-2 N sgRNA qPCR primer targets the shared region of gRNA and sgRNAs, and gRNA qPCR primer targets nsp12. (C) Protein domain enrichment analysis of host factors identified after statistical comparison over no-probe control experiment (i.e. Probe I and Probe II binding). Threshold for statistical significance (P-value < 0.05) is indicated by vertical dashed lines. (D) Validation of RNP capture by silver staining in SARS-CoV-2-infected and uninfected Vero cells. (E) Distribution of peptide length with “on-bead” digestion in SARS-CoV-2-infected and uninfected Vero cells (n = 3 technical replicates). (F) Gene Ontology (GO) Biological Process (BP) term enrichment analysis of 37 host proteins statistically enriched in both Probe I and Probe II experiments.

**Figure S2.**
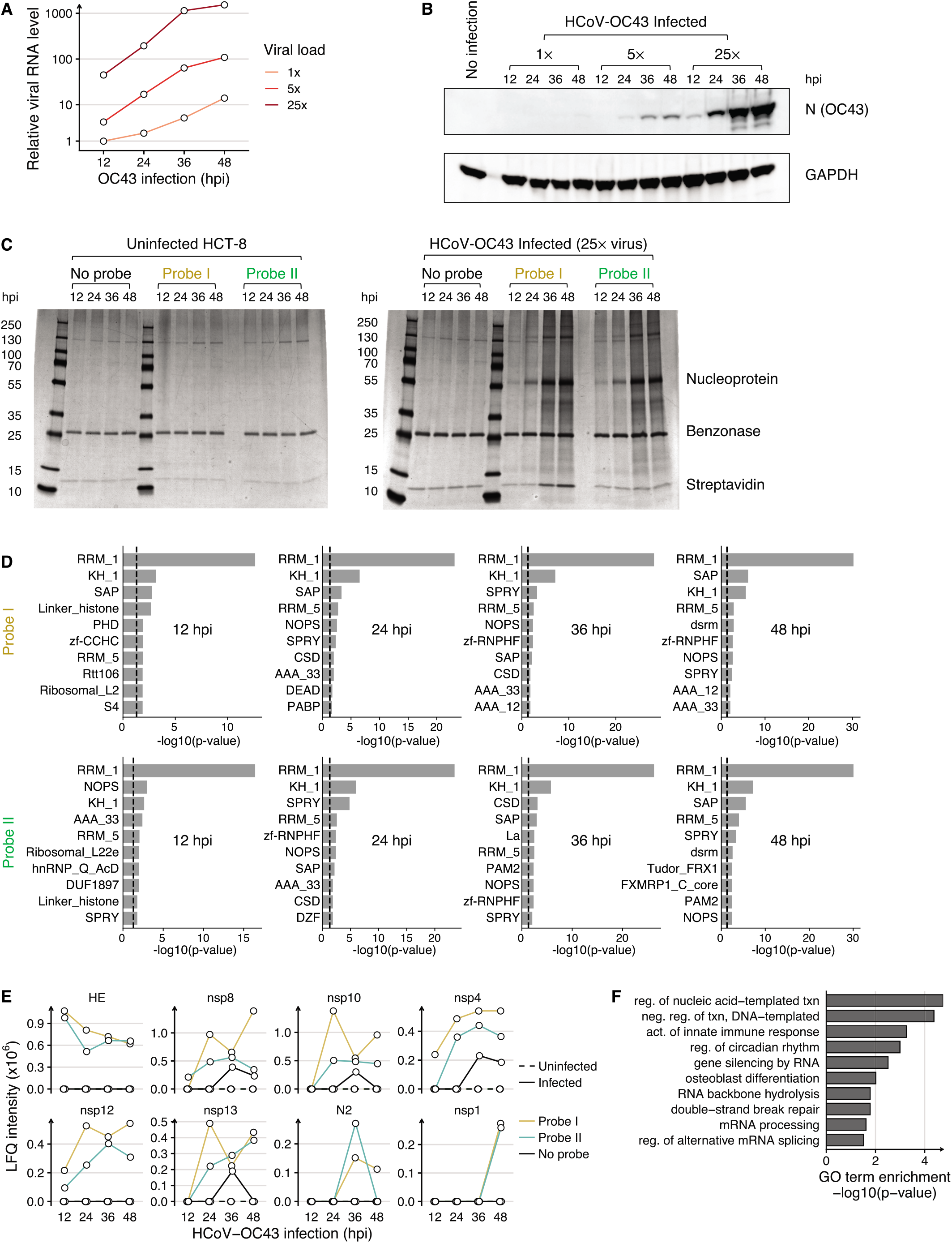
Experimental and statistical validation of RNP capture on HCoV-OC43 RNAs. (A) Relative HCoV-OC43 gRNA (amplicon nsp12) level measured by RT-qPCR and (B) HCoV-OC43 N protein level measured by western blotting for multiple viral loads (1×, 5×, and 25×) across post-infection timepoints of 12, 24, 36, and 48 hours. RNA levels are relative to gRNA levels at 12 hours after 1x viral load infection. (C) Confirmation of RNP capture by silver staining in HCoV-OC43-infected HCT-8 cells. (D) Protein domain enrichment analysis of host factors identified after statistical comparison over no-probe control experiment (i.e. Probe I and Probe II binding) for each post-infection timepoint RNP capture. Threshold for statistical significance (P-value < 0.05) is indicated by vertical dashed lines. (E) LFQ intensity of viral proteins identified in HCoV-OC43 RNP capture at 12, 24, 36, and 48 hours post-infection (hpi) not shown in Figure 2B. (F) Gene Ontology (GO) Biological Process (BP) term enrichment analysis of the 38 host proteins statistically enriched in both OC43 Probe I and Probe II experiments. txn: transcription

**Figure S3.**
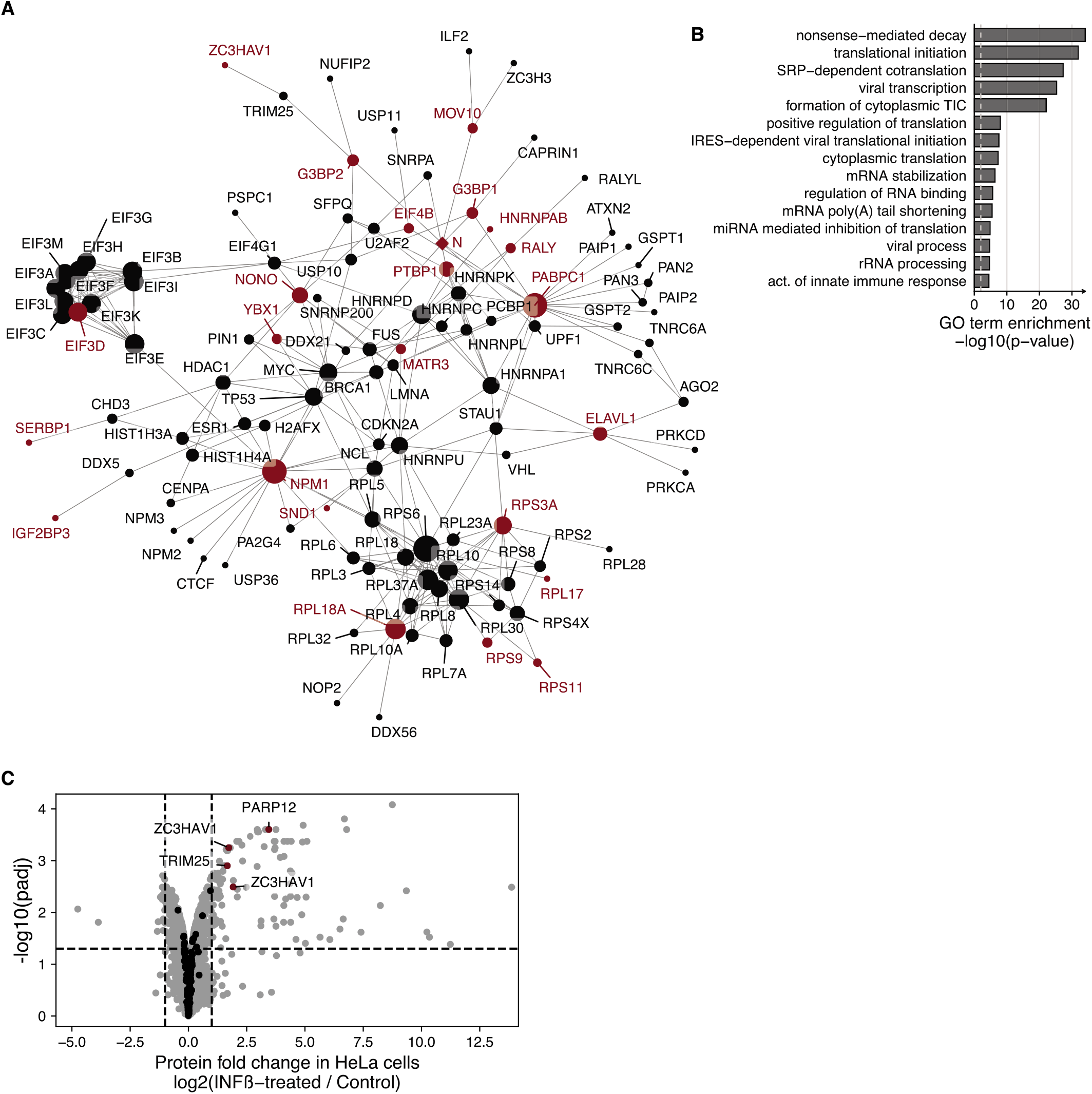
The SARS-CoV-2 RNA interactome in the context of the physical-interaction network and gene-regulatory network. (A) Physical-interaction network of host and viral proteins enriched in both SARS-CoV-2 Probe I and Probe II experiments which are designated in red. Host and viral proteins are marked by circles and diamonds, respectively. The size of the network node is proportional to the number of network edges (i.e. degree) of the node. Largest connected component of the network was chosen for visualization and interpretation. (B) GO BP term enrichment analysis of host proteins retrieved in (A). (C) Differential expression analysis of published proteomic data of HeLa cells after 24 hours of INFß stimulation. Protein-level fold change (FC) of −1 and 1 are marked by vertical dashed lines. 1% FDR is marked by a horizontal dashed line. Host proteins of the SARS-CoV-2 RNA interactome are marked in red if differentially expressed (FDR < 1% and |FC| > 1) or otherwise in black.

**Figure S4.**
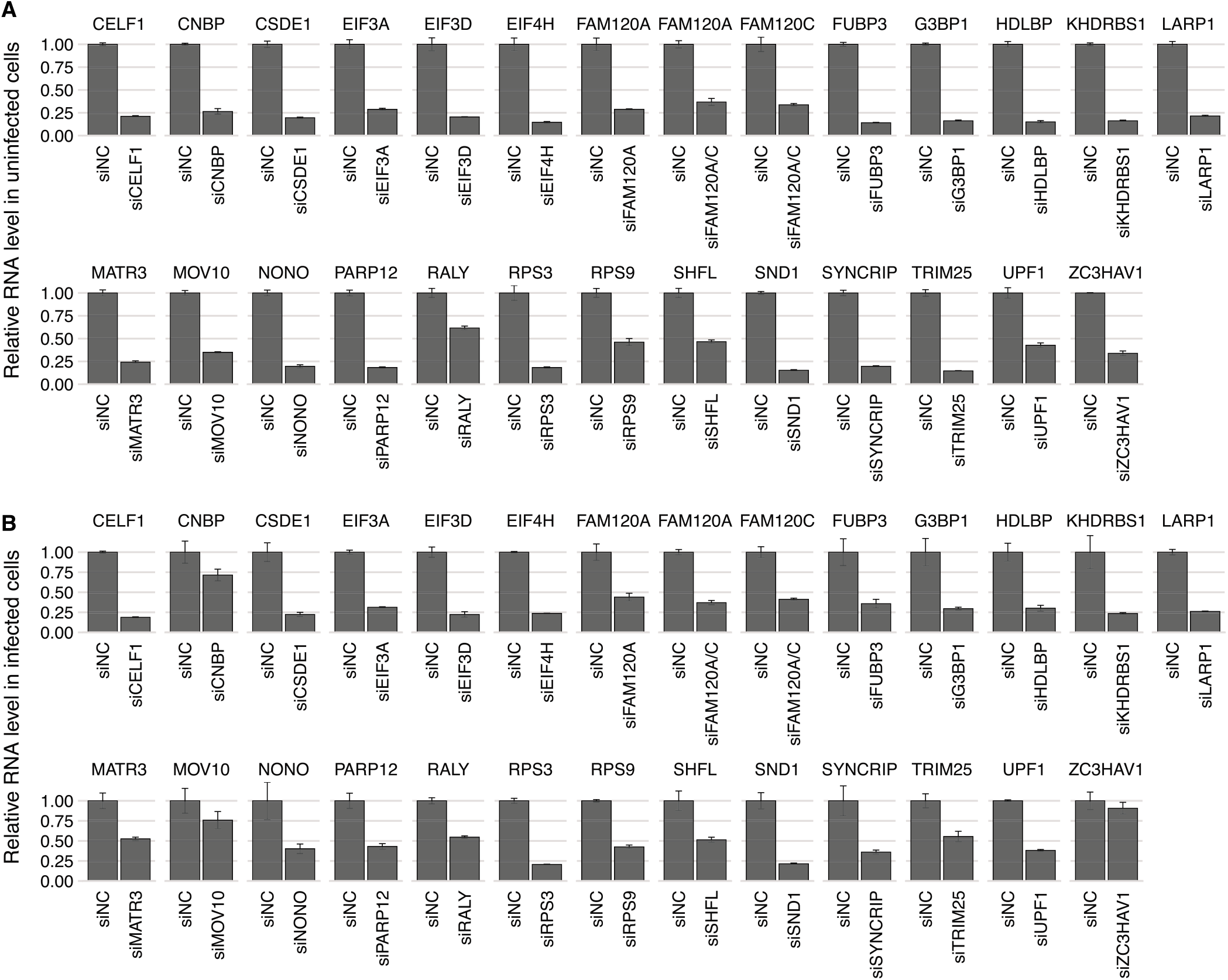
Experimental confirmation of siRNA knockdown in Calu-3 cells. siRNA knockdown confirmation by RT-qPCR in (A) Calu-3 cells and (B) SARS-CoV-2-infected Calu-3 cells 24 hours after infection at 0.05 MOI. Data are represented as mean ± s.e.m. (n□=□3 independent experiments). 18S rRNA was used for normalization.

## Methods

### Preparation of antisense oligonucleotide templates

By scanning the genomic RNAs of SARS-CoV-2 (NCBI RefSeq accession NC_045512.2) and HCoV-OC43 (GenBank accession AY391777.1) from head to tail, partially overlapping 90 nt tiles were enumerated. These tiles were designed to have 30 nt spacing, so adjacent tiles share a subsequence of 60 nt. To avoid ambiguous targeting, tiles were aligned to the human transcriptome (version of Oct 14, 2019) using bowtie 2 (Langmead and Salzberg, 2012) and multi-mapped sequences were discarded. To prepare biotinylated antisense oligonucleotides (ASOs) in bulk, the sequence elements for in vitro transcription (IVT), reverse transcription (RT) and PCR were added to the 90 nt tiles. The T7 promoter (5’-TAA TAC GAC TCA CTA TAG GG-3’) and a pad for RT priming (5’-TGG AAT TCT CGG GTG CCA AGG-3’) were added to the head and tail of each tile, respectively. We grouped ASOs into two sets for each viral genome: “Probe I” targets the unique region of genomic RNA ([265:21553] of NC_045512.2; [:21506] of AY391777.1) and “Probe II” aims at both genomic and subgenomic RNAs ([21562:] of NC_045512.2; [21506:] of AY391777.1). The templates of four ASO groups have distinct PCR primer binding sites on both ends. Accordingly, each ASO set can be selectively amplified from a single mixture. The final ASO templates (167 nt) were prepared via the oligo pool synthesis service of Genscript and stored at −80°C. The ASO templates used in this study are listed in Table S2.

### Mass production of biotin-labeled ASO

ASO templates were amplified using KAPA HiFi HotStart ReadyMix (Roche) and PCR primers for an ASO pool. PCR products were purified by QIAquick PCR purification kit (QIAGEN). RNA intermediates were then transcribed using 5X MEGAscript T7 transcription kit (Invitrogen), and DNA templates were degraded by TURBO DNase (Invitrogen). To clean up enzymes and other reagents, 1.8× reaction volume of AMPure XP (Beckman) was applied and polyethylene glycol was added to be final 20%. The size selection was carried out according to the manufacturer’s protocol. Biotinylated ASOs were synthesized by RevertAid Reverse Transcriptase (Thermo Scientific) and 5’ biotin-TEG primer. RNA intermediates were hydrolyzed at 0.1 M NaOH and neutralized with acetic acid. Finally, ASO purification was performed in the same manner as IVT RNA selection. The primer sequences used for PCR and reverse transcription are listed in Table S6.

### Compilation of proteome databases

The Uniprot reference proteome sets for human (UP000005640, canonical, SwissProt) and African green monkey (Chlorocebus sabaeus; UP000029965, canonical, SwissProt and TrEMBL) were used to identify host proteins in each mass spectrometry experiment (version 03/21/2020) (UniProt Consortium, 2019). The reference proteome set for the Severe acute respiratory syndrome coronavirus 2 (SARS-CoV-2) was manually curated largely based on the NCBI Reference Sequence (NC_045512.2) and related literature of other accessory proteins (e.g. ORF3b, ORF9b and ORF9c). The reference proteome set for the Human coronavirus OC43 (HCoV-OC43) was compiled based on the Uniprot Swiss-Prot proteins for HCoV-OC43 (taxonomy:31631) except for HCoV-OC43 Protein I which was separated into Protein Ia and Protein Ib (or N2) (Vijgen et al., 2005).

### Cell culture, transfection and virus infection

Virus experiments were carried out in accordance with the biosafety guideline by the Korea Centers for Disease Control & Prevention (KCDC). The Institutional Biosafety Committee of Seoul National University (SNUIBC) approved the protocols used in this study, including SNUIBC-R200331-1-1 for BL2 experiments and SNUIBC-200508-1 for BL3 experiments. Vero and HCT-8 cells were maintained in DMEM (Welgene) and RPMI 1640 (Welgene) respectively, both with 1X Antibiotic-Antimycotic (Gibco) and 10% FBS (Gibco) and cultured in CO_2_ incubator with 5% CO_2_ 37 DC. For SARS-CoV-2 infection, 7 ⍰ 10^6^ Vero cells were plated in T-175 flasks 24 hours before infection. Cells were washed with serum-free media and incubated with 5 mL virus-diluted media for 30 minutes at 0.1 MOI, as determined by plaque assay. After infection, virus containing media was replaced with reduced-serum media (2% FBS) and cultured until the harvest. For HCoV-OC43 infection, a similar protocol was used except for incubation temperature lowered to 33 DC. For siRNA transfection, 3.5 ⍰ 10^5^ Calu-3 cells, maintained in DMEM with 1X Antibiotic-Antimycotic and 10% FBS in CO_2_ incubator with 5% CO_2_ 37 □C, were plated in 12 well plate and final 50 nM siRNAs were reverse-transfected using Lipofectamine RNAiMAX (Invitrogen) and ON-TARGETplus SMARTpool siRNAs (Horizon Discovery). Cell viability after siRNA knockdown was measured by splitting 1/100^th^ of cells from uninfected cells, 48 hours after transfection into 96 well plates in triplicates and cell number was measured by MTT assay (Promega) at 4 hours after addition of tetrazolium dye.

### RNA purification and RT-qPCR

For total RNA purification from virus-infected cells, 1 mL TRIzol LS (Invitorgen) were added to media-removed cell monolayers per single well of 12 well plates followed by on-column DNA digestion and purification (Zymo Research). For RNA purification from RNP capture sample, bead-captured RNAs were digested with 100 ng Proteinase K (PCR grade, Roche) and incubated at 37°C for 1 hour, followed by RNA isolation by TRIzol LS with GlycoBlue (Invitrogen). 1~5 μg RNA were reverse-transcribed using RevertAid transcriptase (Thermo Scientific) and random hexamer. qPCR was performed with primer pairs listed in Table S6 and PowerSYBR Green (Applied Biosystems) and analyzed with QuantStudio 5 (Thermo Scientific).

### Modified RNA antisense purification coupled with mass spectrometry (RAP-MS)

Virus infected cells were detached from culture vessels by trypsin and cell pellets were resuspended with ice-cold PBS. 12 mL cell suspensions were dispersed in 150 mm dishes to irradiate 254 nm UV for 2.5 J/cm^2^ using BIO-LINK BLX-254 for SARS-CoV-2 or 0.8 J/cm^2^ using Spectrolinker XL-1500 for HCoV-OC43. UV-crosslinked cells were pelleted by centrifugation and resuspended in TURBO DNase solution (150 Units per flask) and incubated at 37°C for 30 minutes. DNA digested cells were supplemented with equal volume of pre-heated 2X lysis buffer (40 mM Tris-Cl at pH 7.5, 1 M LiCl, 1% LDS, 2 mM EDTA, 10 mM DTT and 8 M urea) and denatured by incubating at 68°C for 30 minutes. Per replicate, cell lysate from 1 flask (T-175) were mixed with 20 μg biotin probe pools (Probe I or Probe II) and hybridized by incubating at 68°C for 30 minutes in final 1 mL volume. Biotin-labeled RNP lysates were supplemented with streptavidin beads (1 mL per replicate, New England Biolabs) and captured by rotating at room temperature overnight. Probe-enriched RNP beads were washed with 1X lysis buffer twice and transferred to fresh tubes, followed by final wash with detergent-free wash buffer (20 mM Tris-Cl at pH 7.5, 0.5 M LiCl, 1 mM EDTA). 1/10^th^ of beads were set aside for assessment of RNA contents by RT-qPCR and another 1/10^th^ of beads were used for silver staining (KOMA biotech). The remaining 8/10^th^ of beads were digested with 100 units of Benzonase nuclease (Millipore) at 37°C for 1 hour. For on-bead peptide digestion, nuclease treated beads were suspended to final 8 M urea and reduced with 10 mM dithiothreitol (DTT), alkylated with 40 mM iodoacetamide (IAA) for 1 hour each at 37°C, and diluted with 50 mM ammonium bicarbonate (ABC) to final 1 M urea. These bead suspensions were supplemented with 300 ng Trypsin (Thermo Scientific, MS grade) and 1 mM CaCl_2_ and digested overnight at 37°C. Peptide solutions were separated from magnetic beads and further processed with HiPPR detergent removal spin columns (Thermo Scientific) and desalted by reverse phase C18 ziptip (Millipore). After the clean up and dry down, the samples were reconstituted with 20 μL of 25 mM ammonium bicarbonate buffer for LC-MS/MS analysis.

### LC-MS/MS analysis

LC-MS/MS analysis was carried out using Orbitrap Fusion Lumos Tribrid MS (Thermo Fisher Scientific) coupled with nanoAcquity UPLC system (Waters). Both analytical capillary columns (100 cm x 75 μm i.d.) and the trap columns (3 cm x 150 μm i.d) were packed in-house with 3 μm Jupiter C18 particles (Phenomenex, Torrance). The long analytical column was placed in a column heater (Analytical Sales and Services) regulated to a temperature of 45°C. The LC flow rate was 300 nL/min and the 100-min linear gradient ranged from 95% solvent A (H_2_O with 0.1% formic acid (Merck)) to 40% solvent B (100% acetonitrile with 0.1% formic acid). Precursor ions were acquired at 120 K resolving power at m/z 200 and the isolation of precursor for MS/MS analysis was performed with a 1.4 Th. Higher-energy collisional dissociation (HCD) with 30% collision energy was used for sequencing with a target value of 1E5 ions determined by automatic gain control. Resolving power for acquired MS2 spectra was set to 30 K at m/z 200 with 120 ms maximum injection time.

Mass spectrometric raw data files were processed for Label-Free Quantification with MaxQuant (version 1.6.15.0) (Cox and Mann, 2008) using the built-in Andromeda search engine (Cox et al., 2011) at default settings with a few exceptions. Briefly, for peptide-spectrum match (PSM) search, cysteine carbamidomethylation was set as fixed modifications, and methionine oxidation and N-terminal acetylation were set as variable modifications. Tolerance for the first and main PSM search were 20 and 4.5 ppm, respectively. Peptides from common contaminant proteins were identified by utilizing the contaminant database provided by MaxQuant. FDR threshold of 1% was used for both the peptide and protein level. The match-between-runs option was enabled with default parameters in the identification step. Finally, LFQ was performed for those with a minimum ratio count of 1.

### Statistical analysis for RNP capture experiment

To identify host and viral proteins that interact with the particular RNA species of interest (e.g. sgRNA or gRNA), we utilized the results from the “bead only” and “probe only” samples as technical backgrounds. Specifically, the “bead only” (or no-probe) experiment in infected cells was used to account for non-specific interactors and biotin-containing carboxylases (e.g. PCCA, ACACA, and ACACB) and determine the set of host and viral proteins that in a broad sense bind to the RNA, which we call Probe I/II “binding” proteins. The probe experiment in uninfected cells (i.e. “probe only”) was then used as the technical background against target RNA-independent interactors and determine the set of host proteins that are enriched for the target RNA, which we call Probe I/II “enriched” proteins.

To accomplish this, we considered the protein spectra count data as a multinomial distribution and applied a statistical test for spectra count enrichment. Specifically, let *N_p_* be the number of identified spectra counts for protein group p from the case experiment (e.g. Probe I experiment in infected cells), and *M_p_* be the respective count number from the control experiment (e.g. no-probe experiment in infected cells). For each protein *i* with *N_i_* ≥ 1, the statistical significance of enrichment is:

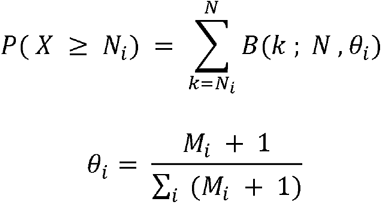

where *N* = ∑_*i*_ *N_i_* is the total spectra count, *θ_i_* is the background probability, and *B*(*k*; *n,p*) is the binomial distribution of *k* successes in *n* trails with success probability *p.* Finally, the Benjamini-Hochberg method was used to adjust the p-values and control the false discovery rate.

### Gene Ontology (GO) enrichment analysis

We conducted enrichment analyses of Gene Ontology (GO) terms (Gene Ontology Consortium, 2001) by means of summarizing the function of tens of host proteins identified in the RNP capture experiment. In general, Fisher’s exact test is used to estimate the statistical significance of the association (i.e. contingency) between a particular GO term and the gene set of interest.

To improve the explanatory power of this analysis, we used the weight01 algorithm (Alexa et al., 2006) from the topGO R package which accounts for the GO graph structure and reduces local dependencies between GO terms. Detailed information of the Gene Ontology was from the GO.db R package (version 3.8.2), and GO gene annotations were from the org.Hs.eg.db R package (version 3.8.2).

### Protein-protein interaction network analysis

We integrated protein-protein interaction data from the BioGRID database (Release 3.5.187) (Stark et al., 2006) and retrieved other proteins that do not necessarily bind to the SARS-CoV-2 RNA but form either transient or stable physical interactions with the host proteins identified from the RNP capture experiments. In detail, we considered only human protein-protein interactions that were (1) found from at least two different types of experiments and (2) reported by at least three publication records which resulted in a total of 65,625 interactions covering 12,143 human proteins. Physical interactions between SARS-CoV-2 proteins and human proteins were by affinity capture and mass spectrometry in SARS-CoV-2 protein expressing cells (Gordon et al., 2020). The network R package and the ggnet2 function of the GGally R package was used for graph visualization.

### Protein domain enrichment analysis

Pfam database (version 33.1) (El-Gebali et al., 2019) was used for protein domain enrichment analysis. Taxon 9606 (human) and Taxon 60711 (green monkey) protein domain annotations were used to analyze RNP capture results of HCoV-OC43 and SARS-CoV-2, respectively. Onesided Fisher’s exact test was applied to estimate the statistical enrichment of a particular protein domain for the specific gene set (e.g. SARS-CoV-2 Probe I binding proteins). We utilized the set of all proteins identified in the RNP capture experiments and all protein domains annotated to those proteins as the statistical background of the enrichment analysis.

### Subcellular localization analysis

To investigate the subcellular localizations of the SARS-CoV-2 interactome, we leveraged the protein subcellular localization information from the Human cell map database v1 (Go et al., 2019). Information from the SAFE algorithm was used primarily but then supplemented by information from the NMF algorithm in case of “no prediction” or localizations.. Localization terms of the NMF algorithm were matched to terms of the SAFE algorithm in general, but few were mapped to the higher term of the SAFE algorithm. For example, the “cell junction” term of the NMF algorithm was merged to the “cell junction, plasma membrane” term of the SAFE algorithm.

## References

Alexa, A., Rahnenführer, J., and Lengauer, T. (2006). Improved scoring of functional groups from gene expression data by decorrelating GO graph structure. Bioinformatics 22, 1600–1607.

Anderson, E.C., Hunt, S.L., and Jackson, R.J. (2007). Internal initiation of translation from the human rhinovirus-2 internal ribosome entry site requires the binding of Unr to two distinct sites on the 5’ untranslated region. J. Gen. Virol. 88, 3043–3052.

Atasheva, S., Frolova, E.I., and Frolov, I. (2014). Interferon-stimulated poly(ADP-Ribose) polymerases are potent inhibitors of cellular translation and virus replication. J. Virol. 88, 2116–2130.

Balinsky, C.A., Schmeisser, H., Wells, A.I., Ganesan, S., Jin, T., Singh, K., and Zoon, K.C. (2017). IRAV (FLJ11286), an Interferon-Stimulated Gene with Antiviral Activity against Dengue Virus, Interacts with MOV10. J. Virol. 91.

Baltz, A.G., Munschauer, M., Schwanhäusser, B., Vasile, A., Murakawa, Y., Schueler, M., Youngs, N., Penfold-Brown, D., Drew, K., Milek, M., et al. (2012). The mRNA-bound proteome and its global occupancy profile on protein-coding transcripts. Mol. Cell 46, 674–690.

Blanco-Melo, D., Nilsson-Payant, B.E., Liu, W.-C., Uhl, S., Hoagland, D., Møller, R., Jordan, T.X., Oishi, K., Panis, M., Sachs, D., et al. (2020). Imbalanced Host Response to SARS-CoV-2 Drives Development of COVID-19. Cell 181, 1036–1045.e9.

Blichenberg, A., Schwanke, B., Rehbein, M., Garner, C.C., Richter, D., and Kindler, S. (1999). Identification of a cis-acting dendritic targeting element in MAP2 mRNAs. J. Neurosci. 19, 8818–8829.

Bojkova, D., Klann, K., Koch, B., Widera, M., Krause, D., Ciesek, S., Cinatl, J., and Münch, C. (2020). Proteomics of SARS-CoV-2-infected host cells reveals therapy targets. Nature 583, 469–472.

Bouhaddou, M., Memon, D., Meyer, B., White, K.M., Rezelj, V.V., Correa Marrero, M., Polacco, B.J., Melnyk, J.E., Ulferts, S., Kaake, R.M., et al. (2020). The Global Phosphorylation Landscape of SARS-CoV-2 Infection. Cell 182, 685–712.e19.

Boussadia, O., Niepmann, M., Créancier, L., Prats, A.-C., Dautry, F., and Jacquemin-Sablon, H. (2003). Unr is required in vivo for efficient initiation of translation from the internal ribosome entry sites of both rhinovirus and poliovirus. J. Virol. 77, 3353–3359.

Bouvet, M., Debarnot, C., Imbert, I., Selisko, B., Snijder, E.J., Canard, B., and Decroly, E. (2010). In vitro reconstitution of SARS-coronavirus mRNA cap methylation. PLoS Pathog. 6, e1000863.

Castello, A., Fischer, B., Eichelbaum, K., Horos, R., Beckmann, B.M., Strein, C., Davey, N.E., Humphreys, D.T., Preiss, T., Steinmetz, L.M., et al. (2012). Insights into RNA biology from an atlas of mammalian mRNA-binding proteins. Cell 149, 1393–1406.

Chapman, E.G., Costantino, D.A., Rabe, J.L., Moon, S.L., Wilusz, J., Nix, J.C., and Kieft, J.S. (2014). The structural basis of pathogenic subgenomic flavivirus RNA (sfRNA) production. Science 344, 307–310.

Chen, C.Z., Shinn, P., Itkin, Z., Eastman, R., Bostwick, R., Rasmussen, L., Huang, R., Shen, M., Hu, X., Wilson, K.M., et al. (2020). Drug Repurposing Screen for Compounds Inhibiting the Cytopathic Effect of SARS-CoV-2. bioRxiv http://dx.doi.org/10.1101/2020.08.18.255877.

Choudhury, N.R., Heikel, G., and Michlewski, G. (2020). TRIM25 and its emerging RNA-binding roles in antiviral defense. WIREs RNA 11, 1.

Chu, C., Zhang, Q.C., da Rocha, S.T., Flynn, R.A., Bharadwaj, M., Calabrese, J.M., Magnuson, T., Heard, E., and Chang, H.Y. (2015). Systematic discovery of Xist RNA binding proteins. Cell 161, 404–416.

Cox, J., and Mann, M. (2008). MaxQuant enables high peptide identification rates, individualized ppb-range mass accuracies and proteome-wide protein quantification. Nat. Biotechnol. 26, 1367–1372.

Cox, J., Neuhauser, N., Michalski, A., Scheltema, R.A., Olsen, J.V., and Mann, M. (2011). Andromeda: a peptide search engine integrated into the MaxQuant environment. J. Proteome Res. 10, 1794–1805.

Dudaronek, J.M., Barber, S.A., and Clements, J.E. (2007). CUGBP1 is required for IFNbeta-mediated induction of dominant-negative CEBPbeta and suppression of SIV replication in macrophages. J. Immunol. 179, 7262–7269.

Egloff, M.-P., Ferron, F., Campanacci, V., Longhi, S., Rancurel, C., Dutartre, H., Snijder, E.J., Gorbalenya, A.E., Cambillau, C., and Canard, B. (2004). The severe acute respiratory syndrome-coronavirus replicative protein nsp9 is a single-stranded RNA-binding subunit unique in the RNA virus world. Proc. Natl. Acad. Sci. U. S. A. 101, 3792–3796.

El-Gebali, S., Mistry, J., Bateman, A., Eddy, S.R., Luciani, A., Potter, S.C., Qureshi, M., Richardson, L.J., Salazar, G.A., Smart, A., et al. (2019). The Pfam protein families database in 2019. Nucleic Acids Res. 47, D427–D432.

Engreitz, J.M., Pandya-Jones, A., McDonel, P., Shishkin, A., Sirokman, K., Surka, C., Kadri, S., Xing, J., Goren, A., Lander, E.S., et al. (2013). The Xist lncRNA exploits three-dimensional genome architecture to spread across the X chromosome. Science 341, 1237973.

Fehr, A.R., Athmer, J., Channappanavar, R., Phillips, J.M., Meyerholz, D.K., and Perlman, S. (2015). The nsp3 macrodomain promotes virulence in mice with coronavirus-induced encephalitis. J. Virol. 89, 1523–1536.

Fehr, A.R., Singh, S.A., Kerr, C.M., Mukai, S., Higashi, H., and Aikawa, M. (2020). The impact of PARPs and ADP-ribosylation on inflammation and host-pathogen interactions. Genes Dev. 34, 341–359.

Finkel, Y., Mizrahi, O., Nachshon, A., Weingarten-Gabbay, S., Yahalom-Ronen, Y., Tamir, H., Achdout, H., Melamed, S., Weiss, S., Israely, T., et al. (2020). The coding capacity of SARS-CoV-2. bioRxiv http://dx.doi.org/10.1101/2020.05.07.082909.

Fung, T.S., and Liu, D.X. (2019). Human Coronavirus: Host-Pathogen Interaction. Annu. Rev. Microbiol. 73, 529–557.

Gack, M.U., Shin, Y.C., Joo, C.-H., Urano, T., Liang, C., Sun, L., Takeuchi, O., Akira, S., Chen, Z., Inoue, S., et al. (2007). TRIM25 RING-finger E3 ubiquitin ligase is essential for RIG-I-mediated antiviral activity. Nature 446, 916–920.

Gene Ontology Consortium (2001). Creating the gene ontology resource: design and implementation. Genome Res. 11, 1425–1433.

Gerstberger, S., Hafner, M., and Tuschl, T. (2014). A census of human RNA-binding proteins. Nat. Rev. Genet. 15, 829–845.

Go, C.D., Knight, J.D.R., Rajasekharan, A., Rathod, B., Hesketh, G.G., Abe, K.T., Youn, J.-Y., Samavarchi-Tehrani, P., Zhang, H., Zhu, L.Y., et al. (2019). A proximity biotinylation map of a human cell. bioRxiv https://doi.org/10.1101/796391.

Goodier, J.L., Pereira, G.C., Cheung, L.E., Rose, R.J., and Kazazian, H.H., Jr (2015). The Broad-Spectrum Antiviral Protein ZAP Restricts Human Retrotransposition. PLoS Genet. 11, e1005252.

Gordon, D.E., Jang, G.M., Bouhaddou, M., Xu, J., Obernier, K., White, K.M., O’Meara, M.J., Rezelj, V.V., Guo, J.Z., Swaney, D.L., et al. (2020). A SARS-CoV-2 protein interaction map reveals targets for drug repurposing. Nature 583, 459–468.

de Groot, R.J., Baker, S.C., Baric, R.S., Brown, C.S., Drosten, C., Enjuanes, L., Fouchier, R.A.M., Galiano, M., Gorbalenya, A.E., Memish, Z.A., et al. (2013). Commentary: Middle East Respiratory Syndrome Coronavirus (MERS-CoV): Announcement of the Coronavirus Study Group. J. Virol. 87, 7790–7792.

Grunewald, M.E., Chen, Y., Kuny, C., Maejima, T., Lease, R., Ferraris, D., Aikawa, M., Sullivan, C.S., Perlman, S., and Fehr, A.R. (2019). The coronavirus macrodomain is required to prevent PARP-mediated inhibition of virus replication and enhancement of IFN expression. PLoS Pathog. 15, e1007756.

Hadjadj, J., Yatim, N., Barnabei, L., Corneau, A., Boussier, J., Smith, N., Péré, H., Charbit, B., Bondet, V., Chenevier-Gobeaux, C., et al. (2020). Impaired type I interferon activity and inflammatory responses in severe COVID-19 patients. Science 369, 718–724.

Hamre, D., and Procknow, J.J. (1966). A new virus isolated from the human respiratory tract. Proc. Soc. Exp. Biol. Med. 121, 190–193.

Hentze, M.W., Castello, A., Schwarzl, T., and Preiss, T. (2018). A brave new world of RNA-binding proteins. Nat. Rev. Mol. Cell Biol. 19, 327–341.

Hong, S., Freeberg, M.A., Han, T., Kamath, A., Yao, Y., Fukuda, T., Suzuki, T., Kim, J.K., and Inoki, K. (2017). LARP1 functions as a molecular switch for mTORC1-mediated translation of an essential class of mRNAs. Elife 6.

Hu, Y., Li, W., Gao, T., Cui, Y., Jin, Y., Li, P., Ma, Q., Liu, X., and Cao, C. (2017). The Severe Acute Respiratory Syndrome Coronavirus Nucleocapsid Inhibits Type I Interferon Production by Interfering with TRIM25-Mediated RIG-I Ubiquitination. J. Virol. 91.

Kerr, C.H., Skinnider, M.A., Andrews, D.D.T., Madero, A.M., Chan, Q.W.T., Stacey, R.G., Stoynov, N., Jan, E., and Foster, L.J. (2020). Dynamic rewiring of the human interactome by interferon signaling. Genome Biol. 21, 140.

Kim, D., Lee, J.-Y., Yang, J.-S., Kim, J.W., Kim, V.N., and Chang, H. (2020a). The Architecture of SARS-CoV-2 Transcriptome. Cell 181, 914–921.e10.

Kim, J.-M., Chung, Y.-S., Jo, H.J., Lee, N.-J., Kim, M.S., Woo, S.H., Park, S., Kim, J.W., Kim, H.M., and Han, M.-G. (2020b). Identification of Coronavirus Isolated from a Patient in Korea with COVID-19. Osong Public Health Res Perspect 11, 3–7.

Konieczny, P., Stepniak-Konieczna, E., and Sobczak, K. (2014). MBNL proteins and their target RNAs, interaction and splicing regulation. Nucleic Acids Res. 42, 10873–10887.

Lahr, R.M., Fonseca, B.D., Ciotti, G.E., Al-Ashtal, H.A., Jia, J.-J., Niklaus, M.R., Blagden, S.P., Alain, T., and Berman, A.J. (2017). La-related protein 1 (LARP1) binds the mRNA cap, blocking eIF4F assembly on TOP mRNAs. Elife 6.

Lai, M.M.C., and Cavanagh, D. (1997). The Molecular Biology of Coronaviruses. In Advances in Virus Research, K. Maramorosch, F.A. Murphy, and A.J. Shatkin, eds. (Academic Press), pp. 1–100.

Lai, M.M., and Stohlman, S.A. (1981). Comparative analysis of RNA genomes of mouse hepatitis viruses. J. Virol. 38, 661–670.

Lan, T.C.T., Allan, M.F., Malsick, L.E., Khandwala, S., Nyeo, S.S.Y., Bathe, M., Griffiths, A., and Rouskin, S. (2020). Structure of the full SARS-CoV-2 RNA genome in infected cells. bioRxiv http://dx.doi.org/10.1101/2020.06.29.178343.

Langmead, B., and Salzberg, S.L. (2012). Fast gapped-read alignment with Bowtie 2. Nat. Methods 9, 357–359.

Lee, F.C.Y., and Ule, J. (2018). Advances in CLIP Technologies for Studies of Protein-RNA Interactions. Mol. Cell 69, 354–369.

Lee, A.S., Kranzusch, P.J., Doudna, J.A., and Cate, J.H.D. (2016). eIF3d is an mRNA cap-binding protein that is required for specialized translation initiation. Nature 536, 96–99.

Lee, K.-M., Chen, C.-J., and Shih, S.-R. (2017). Regulation Mechanisms of Viral IRES-Driven Translation. Trends Microbiol. 25, 546–561.

Li, L., Zhao, H., Liu, P., Li, C., Quanquin, N., Ji, X., Sun, N., Du, P., Qin, C.-F., Lu, N., et al. (2018). PARP12 suppresses Zika virus infection through PARP-dependent degradation of NS1 and NS3 viral proteins. Sci. Signal. 11.

Manokaran, G., Finol, E., Wang, C., Gunaratne, J., Bahl, J., Ong, E.Z., Tan, H.C., Sessions, O.M., Ward, A.M., Gubler, D.J., et al. (2015). Dengue subgenomic RNA binds TRIM25 to inhibit interferon expression for epidemiological fitness. Science 350, 217–221.

Masutani, M., Sonenberg, N., Yokoyama, S., and Imataka, H. (2007). Reconstitution reveals the functional core of mammalian eIF3. EMBO J. 26, 3373–3383.

McHugh, C.A., and Guttman, M. (2018). RAP-MS: A Method to Identify Proteins that Interact Directly with a Specific RNA Molecule in Cells. Methods Mol. Biol. 1649, 473–488.

McHugh, C.A., Chen, C.-K., Chow, A., Surka, C.F., Tran, C., McDonel, P., Pandya-Jones, A., Blanco, M., Burghard, C., Moradian, A., et al. (2015). The Xist lncRNA interacts directly with SHARP to silence transcription through HDAC3. Nature 521, 232–236.

Miknis, Z.J., Donaldson, E.F., Umland, T.C., Rimmer, R.A., Baric, R.S., and Schultz, L.W. (2009). Severe acute respiratory syndrome coronavirus nsp9 dimerization is essential for efficient viral growth. J. Virol. 83, 3007–3018.

Minajigi, A., Froberg, J., Wei, C., Sunwoo, H., Kesner, B., Colognori, D., Lessing, D., Payer, B., Boukhali, M., Haas, W., et al. (2015). Chromosomes. A comprehensive Xist interactome reveals cohesin repulsion and an RNA-directed chromosome conformation. Science 349.

Mukherjee, J., Hermesh, O., Eliscovich, C., Nalpas, N., Franz-Wachtel, M., Maček, B., and Jansen, R.-P. (2019). β-Actin mRNA interactome mapping by proximity biotinylation. Proc. Natl. Acad. Sci. U. S. A. 116, 12863–12872.

Narayanan, K., Huang, C., Lokugamage, K., Kamitani, W., Ikegami, T., Tseng, C.-T.K., and Makino, S. (2008). Severe acute respiratory syndrome coronavirus nsp1 suppresses host gene expression, including that of type I interferon, in infected cells. J. Virol. 82, 4471–4479.

Ooi, Y.S., Majzoub, K., Flynn, R.A., Mata, M.A., Diep, J., Li, J.K., van Buuren, N., Rumachik, N., Johnson, A.G., Puschnik, A.S., et al. (2019). An RNA-centric dissection of host complexes controlling flavivirus infection. Nat Microbiol 4, 2369–2382.

Peiris, J.S.M., Lai, S.T., Poon, L.L.M., Guan, Y., Yam, L.Y.C., Lim, W., Nicholls, J., Yee, W.K.S., Yan, W.W., Cheung, M.T., et al. (2003). Coronavirus as a possible cause of severe acute respiratory syndrome. Lancet 361, 1319–1325.

Perlman, S., and Netland, J. (2009). Coronaviruses post-SARS: update on replication and pathogenesis. Nature Reviews Microbiology 7, 439–450.

Philippe, L., Vasseur, J.-J., Debart, F., and Thoreen, C.C. (2018). La-related protein 1 (LARP1) repression of TOP mRNA translation is mediated through its cap-binding domain and controlled by an adjacent regulatory region. Nucleic Acids Res. 46, 1457–1469.

Ramanathan, M., Porter, D.F., and Khavari, P.A. (2019). Methods to study RNA–protein interactions. Nat. Methods 16, 225–234.

Rogers, G.W., Jr, Richter, N.J., Lima, W.F., and Merrick, W.C. (2001). Modulation of the helicase activity of eIF4A by eIF4B, eIF4H, and eIF4F. J. Biol. Chem. 276, 30914–30922.

Roth, A., and Diederichs, S. (2015). Molecular biology: Rap and chirp about X inactivation. Nature 521, 170–171.

Sa Ribero, M., Jouvenet, N., Dreux, M., and Nisole, S. (2020). Interplay between SARS-CoV-2 and the type I interferon response. PLoS Pathog. 16, e1008737.

Shaw, A.E., Hughes, J., Gu, Q., Behdenna, A., Singer, J.B., Dennis, T., Orton, R.J., Varela, M., Gifford, R.J., Wilson, S.J., et al. (2017). Fundamental properties of the mammalian innate immune system revealed by multispecies comparison of type I interferon responses. PLoS Biol. 15, e2004086.

Sims, A.C., Burkett, S.E., Yount, B., and Pickles, R.J. (2008). SARS-CoV replication and pathogenesis in an in vitro model of the human conducting airway epithelium. Virus Res. 133, 33–44.

Snijder, E.J., Decroly, E., and Ziebuhr, J. (2016). The Nonstructural Proteins Directing Coronavirus RNA Synthesis and Processing. Adv. Virus Res. 96, 59–126.

Sola, I., Almazán, F., Zúñiga, S., and Enjuanes, L. (2015). Continuous and Discontinuous RNA Synthesis in Coronaviruses. Annu. Rev. Virol. 2, 265–288.

Stark, C., Breitkreutz, B.-J., Reguly, T., Boucher, L., Breitkreutz, A., and Tyers, M. (2006). BioGRID: a general repository for interaction datasets. Nucleic Acids Res. 34, D535–D539.

St-Germain, J.R., Astori, A., and Samavarchi-Tehrani, P. (2020). A SARS-CoV-2 BioID-based virus-host membrane protein interactome and virus peptide compendium: new proteomics resources for COVID-19 research. bioRxiv https://doi.org/10.1101/2020.08.28.269175.

Sutton, G., Fry, E., Carter, L., Sainsbury, S., Walter, T., Nettleship, J., Berrow, N., Owens, R., Gilbert, R., Davidson, A., et al. (2004). The nsp9 replicase protein of SARS-coronavirus, structure and functional insights. Structure 12, 341–353.

Suzuki, Y., Chin, W.-X., Han, Q. ‘en, Ichiyama, K., Lee, C.H., Eyo, Z.W., Ebina, H., Takahashi, H., Takahashi, C., Tan, B.H., et al. (2016). Characterization of RyDEN (C19orf66) as an Interferon-Stimulated Cellular Inhibitor against Dengue Virus Replication. PLoS Pathog. 12, e1005357.

Takata, M.A., Gonçalves-Carneiro, D., Zang, T.M., Soll, S.J., York, A., Blanco-Melo, D., and Bieniasz, P.D. (2017). CG dinucleotide suppression enables antiviral defence targeting non-self RNA. Nature 550, 124–127.

Tauber, D., Tauber, G., Khong, A., Van Treeck, B., Pelletier, J., and Parker, R. (2020). Modulation of RNA Condensation by the DEAD-Box Protein eIF4A. Cell 180, 411–426.e16.

Thoms, M., Buschauer, R., Ameismeier, M., Koepke, L., Denk, T., Hirschenberger, M., Kratzat, H., Hayn, M., Mackens-Kiani, T., Cheng, J., et al. (2020). Structural basis for translational shutdown and immune evasion by the Nsp1 protein of SARS-CoV-2. Science.

Ule, J., Hwang, H.-W., and Darnell, R.B. (2018). The Future of Cross-Linking and Immunoprecipitation (CLIP). Cold Spring Harb. Perspect. Biol. 10.

UniProt Consortium (2019). UniProt: a worldwide hub of protein knowledge. Nucleic Acids Res. 47, D506–D515.

Van Nostrand, E.L., Freese, P., Pratt, G.A., Wang, X., Wei, X., Xiao, R., Blue, S.M., Chen, J.-Y., Cody, N.A.L., Dominguez, D., et al. (2020). A large-scale binding and functional map of human RNA-binding proteins. Nature 583, 711–719.

Vijgen, L., Keyaerts, E., Moës, E., Thoelen, I., Wollants, E., Lemey, P., Vandamme, A.-M., and Van Ranst, M. (2005). Complete genomic sequence of human coronavirus OC43: molecular clock analysis suggests a relatively recent zoonotic coronavirus transmission event. J. Virol. 79, 1595–1604.

Wang, X., Xuan, Y., Han, Y., Ding, X., Ye, K., Yang, F., Gao, P., Goff, S.P., and Gao, G. (2019). Regulation of HIV-1 Gag-Pol Expression by Shiftless, an Inhibitor of Programmed −1 Ribosomal Frameshifting. Cell 176, 625–635.e14.

Wei, J., Alfajaro, M., Hanna, R., DeWeirdt, P., and Strine, M. (2020). Genome-wide CRISPR screen reveals host genes that regulate SARS-CoV-2 infection. bioRxiv https://doi.org/10.1101/2020.06.16.155101.

Welsby, I., Hutin, D., Gueydan, C., Kruys, V., Rongvaux, A., and Leo, O. (2014). PARP12, an interferon-stimulated gene involved in the control of protein translation and inflammation. J. Biol. Chem. 289, 26642–26657.

Woo, P.C.Y., Lau, S.K.P., Lam, C.S.F., Lau, C.C.Y., Tsang, A.K.L., Lau, J.H.N., Bai, R., Teng, J.L.L., Tsang, C.C.C., Wang, M., et al. (2012). Discovery of seven novel Mammalian and avian coronaviruses in the genus deltacoronavirus supports bat coronaviruses as the gene source of alphacoronavirus and betacoronavirus and avian coronaviruses as the gene source of gammacoronavirus and deltacoronavirus. J. Virol. 86, 3995–4008.

Zhou, P., Yang, X.-L., Wang, X.-G., Hu, B., Zhang, L., Zhang, W., Si, H.-R., Zhu, Y., Li, B., Huang, C.-L., et al. (2020). A pneumonia outbreak associated with a new coronavirus of probable bat origin. Nature 579, 270–273.

